# Agewise mapping of genomic oxidative DNA modification demonstrates oxidative-driven reprogramming of pro-longevity genes

**DOI:** 10.1101/2020.02.17.951582

**Authors:** Lynn Htet Htet Aung, Yin Wang, Zi-Qian Liu, Zhe Li, Zhongjie Yu, Xiatian Chen, Jinning Gao, Peipei Shan, Zhixia Zhou, Peifeng Li

## Abstract

The accumulation of unrepaired oxidatively damaged DNA can influence both the rate of ageing and life expectancy of an organism. Mapping oxidative DNA damage sites at whole-genome scale will help us to recognize the damage-prone sequence and genomic feature information, which is fundamental for ageing research. Here, we developed an algorithm to map the whole-genome oxidative DNA damage at single-base resolution using Single-Molecule Real-Time (SMRT) sequencing technology. We sequenced the genomic oxidative DNA damage landscape of *C. elegans* at different age periods to decipher the potential impact of genomic DNA oxidation on physiological ageing. We observed an age-specific pattern of oxidative modification in terms of motifs, chromosomal distribution, and genomic features. Integrating with RNA-Seq data, we demonstrated that oxidative modification in promoter regions was negatively associated with the expression of pro-longevity genes, denoting that oxidative modification in pro-longevity genes may exert epigenetic potential and thus affect lifespan determination. Together, our study opens up a new field for exploration of “oxigenetics,” that focuses on the mechanisms of redox-mediated ageing.

**Summary:**

- We developed an algorithm to map the oxidative DNA damage at single-base resolution.
- Oxidative DNA damage landscape in *C. elegans* illustrated an age-specific pattern in terms of motifs, chromosomal distribution, and genomic features.
- Oxidative modification in older worms occurred higher frequency at the sex chromosome, with the preference for promoter and exon regions.
- Oxidative modification in promoter regions of pro-longevity genes was negatively associated with their expression, suggesting the oxidative-driven transcript reprogramming of pro-longevity genes in physiological ageing.

## Introduction

Ageing is characterized by a series of changes at both cellular and molecular levels, including senescence, telomere shortening, and changes in gene expression, that lead to impaired functions and increased vulnerability to death (Collado et al.; Lopez-Otin et al., 2013). Although the molecular basis of ageing has been intensively studied for decades, few promising conclusions regarding the underlying mechanisms and anti-ageing treatments, have been reached. Recent studies in multiple organ systems suggest that the rate of ageing is determined by both genetic and environmental factors (Fallin and Matteini, 2009; Rando and Chang, 2012; Speakman et al., 2015). One of the well-established theories for ageing is that of the accumulation of damaged DNAs, which sees aging as a consequence of the unrepaired accumulation of naturally occurring oxidative DNAs (Cadet and Davies, 2017; Cooke and Evans, 2007; Liochev, 2013; Pérez et al., 2009). However, some studies have shown that oxidative DNA damage plays a subtle role in normal ageing; rather it influences the pathological phenotypes experiencing chronic oxidative stress (Salmon et al., 2010; Zawia et al., 2009). Hence, studying the genetic basis of longevity and healthy ageing in model organisms will provide important biological insights to understand the impact of oxidative DNA damage on physiological ageing.

Oxidative DNA damage is an inevitable consequence of cellular metabolism, with gradually increased levels in organismal ageing due to age-associated deterioration in repair mechanisms (Collado et al.; Cooke et al., 2003; Lopez-Otin et al., 2013). Among more than 20 identified oxidative DNA base lesions, 8-oxo-2’deoxyguanosine (8-OHdG) is the most common and well-studied biomarker for oxidative DNA damage (Cooke et al., 2003; Thompson, 2004; Wallace, 2002). 8-OHdG can be quantified in tissue as well as in body fluids, such as urine, blood, and cerebrospinal fluid by HPLC-based analysis (Lagadu et al., 2010; Weimann et al., 2002), and specific antibody assays (Park et al., 1992). However, identification of specific genomic locations or oxidized genes are yet to be achieved. Previously, we identified that oxidative modification in miR-184 associates with the 3’ UTRs of Bcl-xL and Bcl-w that are not its native targets. The mismatch of oxidized miR-184 with Bcl-xL and Bcl-w initiates apoptosis in cardiomyocytes (Wang et al., 2015). This study highlights the critical need for the localization of oxidative modification in the genome. Therefore, for an in-depth analysis of ageing-associated oxidative DNA damage and to understand the molecular mechanisms underlying ageing, it is imperative to map the oxidized bases in a whole-genome scale, which is fundamental for identification of damage-prone and inefficiently repaired regions of DNA. Here, we pursued the whole genomic 8-OHdG landscape in *C. elegans* at different age periods to craft an oxidative DNA damage map of physiological ageing

Recently, several methods for whole-genome oxidative DNA mapping have been reported (Ding et al., 2017; Poetsch et al., 2018; Wu et al., 2018; Yoshihara et al., 2014); nonetheless, all these studies faced the limitation of zooming into single-base resolution and accurately sequence high-GC-rich regions. Single-Molecule Real-Time (SMRT) sequencing (PacBio) can achieve higher accuracy for GC-rich regions, where 8-OHdG modification frequently takes place (Wu et al., 2018), and is able to provide single-base resolution with longer subread lengths. Moreover, SMRT is highly efficient by the simultaneous detection of multiple forms of damaged DNA bases (Bickhart et al., 2017; Clark et al., 2011; Flusberg et al., 2010; Rasko et al., 2011). Recently, SMRT technology was updated to its second-generation Sequel I and II platforms, which perform ∼8 to 8^2^ times more efficiently than its predecessor (RSII), depending on the sequencing mode and chemistry (Ardui et al., 2018; Wenger et al., 2019). However, the feasibility of using Sequel in the detection of 8-OHdG modification has not yet been verified. Therefore, we attempted to map the genome-wide architecture of oxidative modification using SMRT Sequel platform.

Here, for the first time, we report the whole genomic oxidative landscape of *C. elegans* at different age groups at single base-pair resolution. Furthermore, we identified several age-specific 8-OHdG modified genes that are involved in developmental and biosynthesis pathways. We observed that the effects of oxidative modification in longevity regulating genes can be age-specific, and that oxidative modification can epigenetically reprogram the expression of pro-longevity genes, which eventually determine life-as well as health-span. Together, our study brings critical insights into the genetic mechanism of ageing mediated by oxidative DNA damage, which we believe is a whole new field of exploration for personalized medicine that we would like to term as “oxigenetics.”

## Results

### SMRT Sequel platform is suitable for the detection of 8-OHdG modification

Previous study has shown that oxidative DNA damages can be detected in SMRT sequencing by their specific polymerase kinetic signatures (Clark et al., 2011). To calibrate the kinetic signature of the 8-OHdG modification in the Sequel platform, we sequenced the synthetic oligonucleotide containing two 8-OHdGs (Fig. S1A and B, Table S1). We measured each base position (n=465) times in average and determined the IPD ratio and coverage of each base position as described (Schadt et al., 2013). The statistically significant difference between IPD value at each position with the in-silico control is determined by the modification score (−log_10_ *p*-value). The putative 8-OHdG events were detected at positions 34 and 57 of the synthetic oligonucleotides with strict filtration criteria determined by IPD ratio, modification score, and coverage (Fig. S1C and D) and the true positive and false positive detection rates of our approach were analysed (Fig. S1D). Compared to previously established protocol (Clark et al., 2011), we improved the analysis algorithm in two aspects: (i) to minimize the false positive rate, we included not only the IPD ratio, but also the statistically significant modification score (*FDR* < 0.05) to our threshold for identification of 8-OHdG signal, and (ii) to reduce the bias of undetected surrounding base modification, we included the IPD ratio value of +1bp and -1bp to our filtration criteria.

At the whole-genome level, we sequenced 8-OHdG modification sites in gDNA extracted from the pcDNA3.1 plasmid treated with different concentrations of H_2_O_2_ and compared the number of damage sites. We found an increase in 8-OHdG-modified sites along with H_2_O_2_ concentration, but when we treated the sample with Ogg1, which can specifically cleave 8-OHdG (van der Kemp et al., 1996), a concomitant decrease in 8-OHdG were observed by both SMRT sequencing and dot-blot assay (Fig. S2A and B). The presence of 8-OHdG in different samples was confirmed by HPLC-MS/MS (Fig. S2C). Although 8-OHdG was thought to have variable motifs (Tang et al., 2019), our data indicated that 8-OHdG occurred in higher frequency next to the downstream “GG/GC” sequence in the plasmid genome (Fig. S2D and E). Our data showed that 8-OHdG could influence the IPD value of surrounding bases (Fig. S2D), in particular an adjacent adenosine “A” (Fig. S3). Therefore, to certify that the signal in adjacent “A” was due to the influence of the surrounding 8-OHdG modification, we constructed a point mutation to an 8-OHdG signal-positive “G” site adjacent to an “A.” As anticipated, no modification signal was detected in the mutated sample (Fig. S3), corroborating that the signal was due to the effects of 8-OHdG modification. Nevertheless, given that the IPD value at individual positions can still be influenced by surrounding modifications, we excluded the “G” within +6bp and -6bp next to 6mA or 4mC modification which are the commonly detected genomic modification, from our analyses.

### The repertoire of oxidative modifications in *C. elegans* genome displays age-specific patterns

For the next level of complexity, we attempted to understand the genomic DNA oxidation landscape and its association with ageing in *C.elegans*, one of the widely accepted model organisms for ageing studies. To this end, we sequenced unamplified gDNA extracted from the 1-, 10-, and 20-day-old *C. elegans* with above 100-fold genome coverage (Fig. 1A, Table S1). The experimental workflow and the number of biological replicates used in this study were summarized in Fig. S4 and Fig. 1A. We observed that oxidatively modified sites account for 0.01 %, 0.45 %, and 0.15 % of the whole genome in D1, D10, and D20, respectively (Fig. 1B and Fig. 2A). We then profiled the density of 8-OHdG sites normalized to total guanines in the *C. elegans* genome, the relative frequency of 8-OHdG (8-OHdG/ G) in D10 was 3-fold higher than that of D20 and 10-fold higher than that of D1 (Fig. 1C), with an overlap of 0.009% of 8-OHdG sites among all age groups (Fig. 1D). Circos plots displayed the global distribution of 8-OHdG across the *C. elegans* genome (Fig. 1E). On motif analysis, TGGGT was the most enriched motif in the D1 group, while TGGAT was the most frequent motif in both D10 and D20 groups (Fig. 1F, Table S2). These motifs were very similar to the 8-OHdG motifs of (GGGGT) *S. cerevisiae* (Wu et al., 2018) but vastly different from the 6mA motifs of (AGAA and GAGG) *C. elegans* (Greer et al., 2015a) and (GGAAT) human glioblastoma cell lines (Poetsch et al., 2018), indicating that our identified motifs are specific and predominant in *C. elegans* 8-OHdG modification. The frequency of 3’ or 5’ base next to 8-OHdG is highly variable between young and older worms. In young worms, the frequency of each base 5’ to 8-OHdG was 30% G, 30% T, 30% A and 10% C, and that of each base 3’ to 8-OHdG was 40% G, 30% T, 15% A and 15% C; however, for the older worms, the highest frequency of G (40%) was observed 5’ to 8-OHdG, and A (30%) achieved the highest frequency 3’ to 8-OHdG. The motif identified in our study is consistent with findings in mouse and yeast genome, where there is a high abundance of GG dimmer in regions containing 8-OHdG sites.

**Figure 1.**
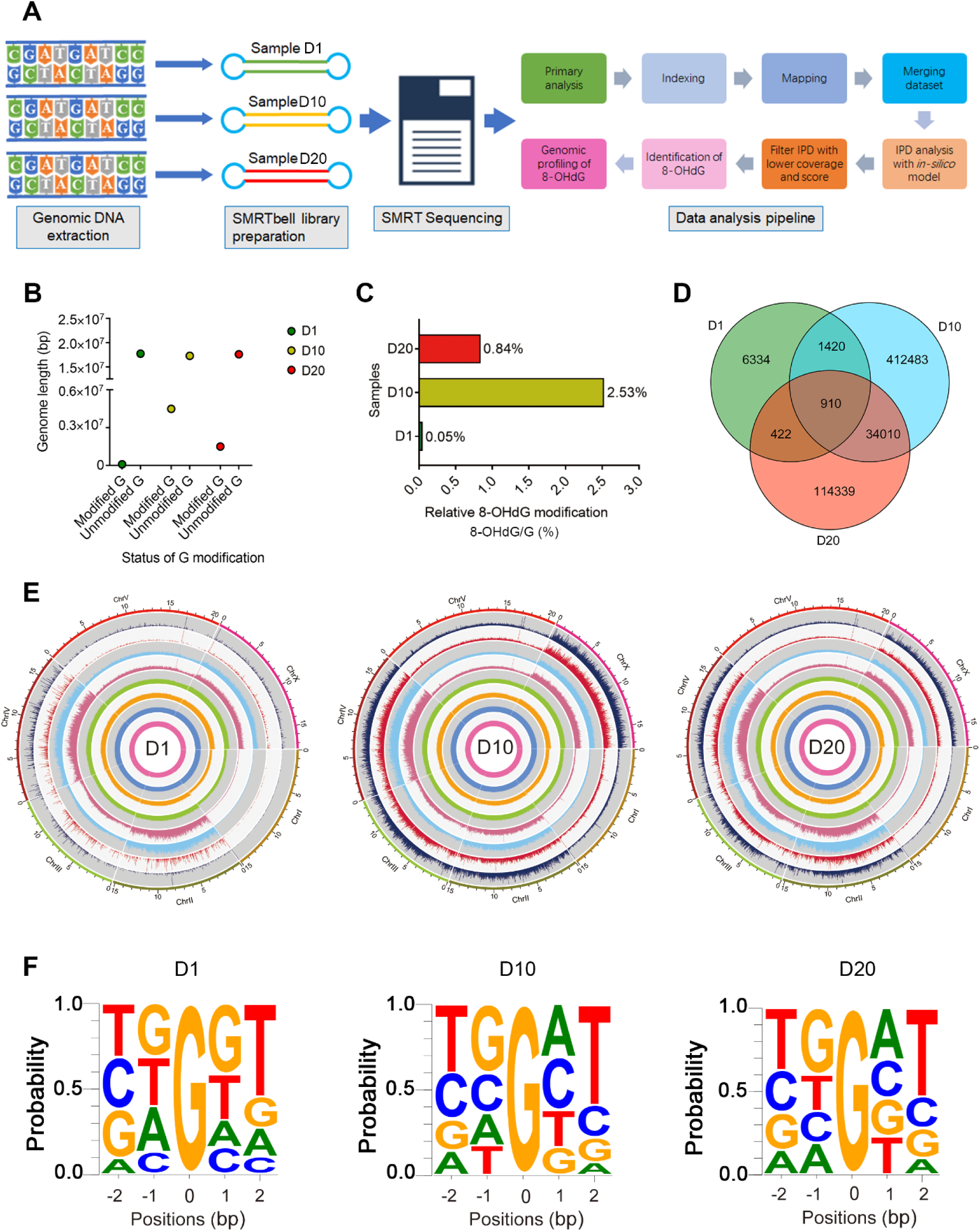
The repertoire of oxidative modifications in *C. elegans* genome displays age-specific patterns. **A**. The workflow of identification of 8-OHdG in the whole-genome level using SMRT sequencing. **B**. The total number of modified and unmodified G-bases observed in different aged groups of *C. elegans*. **C**. Relative frequency of 8-OHdG among different age groups (8-OHdG/G %). **D.** Overlap of 8-OHdG modified sites in 1-day-, 10-day- and 20-day-old *C.elegans*. **E**. Circos plots of 8-OHdG modification profiles of different aged *C. elegan*s. From inner to outer circles: 1^st^ and 2^nd^ indicate IPD ratios, 3^rd^ and 4^th^ represent the coverage of each position detected, 5^th^ and 6^th^ indicate the modification score (−log_10_ *p*-value) for each position tested, 7^th^ and 8^th^ denote 8-OHdG modification for the forward and reversed strands, 9^th^ indicates chromosomes respectively, across 1-day-old, 10-day-old and 20-day-old *C. elegans*. **F**. Motif analysis based on consensus sequences next to 8-OHdG sites.

**Figure 2.**
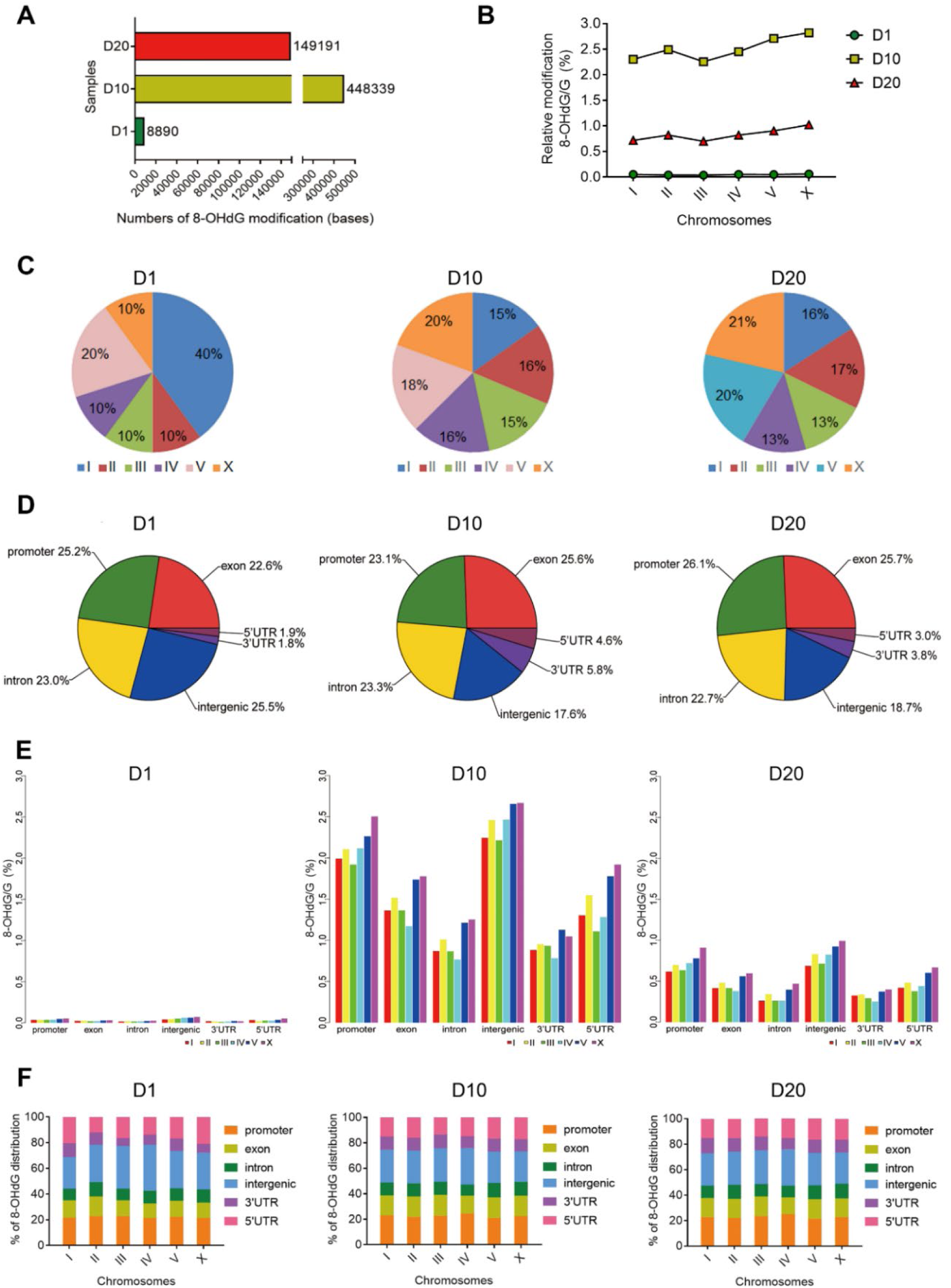
The trends and characteristics of 8-OHdG modification show heterogeneity among different age groups. **A**. The total number of 8-OHdG bases detected in different samples. **B**. The relative frequency of 8-OHdG/G across different chromosomes. To avoid the influence of number of G occupied in each chromosome on the rate of 8-OHdG, the 8-OHdG count was normalized by the total number of G in each chromosome. **C** shows the dynamics of 8-OHdG modifications across different chromosomes. **D**. Pie chart of the distribution of 8-OHdG sites in different genomic regions. We analyzed 8-OHdG distribution in genomic regions divided into promoters, 5’UTR, 3’UTR, exon and intron regions. **E and F**. 8-OHdG modifications detected in different genomic regions of each chromosome. **E** shows the relative frequency, and **F** shows the proportion of 8-OHdG modifications per each chromosome.

### The trends and characteristics of 8-OHdG modification show heterogeneity among different age groups

To understand whether oxidative modification has any chromosomal dominant, we analyzed the frequencies of 8-OHdG across all six chromosomes. The chromosomal distribution of 8-OHdG showed a different trend between young and older worms (Fig. 2B). Of the 6 chromosomes, ChX had the highest frequency of 8-OHdG modification in older worms (D10 and D20), whereas ChI was the lowest; conversely, the trend in young worms was completely opposite to the one observed in older worms (Fig. 2C), demonstrating that the chromosomal distribution pattern of 8-OHdG in *C. elegans* genome is age-specific.

To determine putative biological functions of 8-OHdG modification, we examined 8-OHdG distribution in genomic features divided into gene bodies, including exon, intron, promoters, 5’UTR, 3’UTR regions, and intergenic regions. The highest % of modification for D1 is at the intergenic (non-coding) regions; however, for D10 and D20, the highest % is seen at exon (coding) regions (Fig. 2D). The dynamics of 8-OHdG distribution in different genomic regions of each chromosome show a similar trend for all age groups (Fig. 2E and F).

To investigate whether 8-OHdG modification shows sequence preference for different genomic regions, we performed motif analysis using the sequence information of -2bp and +2bp next to the modification. As described in Figure S5A-C, TTGGT is the most frequent motif found in promoter, 3’UTR, 5’UTR, and TF-binding regions of D1, whereas TTGAT is the most frequent one in D10 and D20 groups. However, for exon regions, TTGAT is the most frequent motif for all age groups. Between D10 and D20, the 8-OHdG motifs for gene body, exon, promoter and 3’UTR are similar but those for 5’UTR and TF-binding regions are different. Overall, although 8-OHdG modification sites indicated significant different motifs between young and older worms, it did not show significant different sequence preferences among different genomic features except TF-binding regions of older worms.

### Oxidative modification patterns change with ageing

Upon analyzing the 8-OHdG modification within the coding regions, we found that 11% (2,124), 92% (18,571), and 72% (14,579) of annotated genes in 1-day-old, 10-day-old, and 20-day-old respectively have at least one 8-OHdG sites. Approximately 9% (1,857) of oxidatively modified genes were overlapped among three age groups, while 60% (12,242) were overlapped between D10 and D20 groups (Fig. 3A and B, and Table S3). The trend of 8-OHdG modified genes across different chromosomes is significantly different between young and older worms; for 1-day old group, the highest number of oxidatively modified genes are found in ChIII, while in 10- and 20-day-old group, the highest number of oxidatively modified genes were in ChV (Fig. 3B and C, and Fig. S6A-C). To understand whether oxidative DNA damage occurred more frequently in coding regions of specific chromosomes, we analyzed the frequency of 8-OHdG (+) genes relative to the total number of genes occupied in each chromosome (Fig. 3D). The highest 8-OHdG (+) genes occupancy was found in ChX for all age groups (Fig. 3D). In D1, the 8-OHdG (+) genes occupancies in ChI and ChIII were higher than that of ChII, while in the older groups (D10 and D20), those in ChI and ChIII were lower than that of ChII (Fig. 3D). These data denoted a different trend of chromosomal 8-OHdG (+) gene distribution among different age groups.

**Figure 3.**
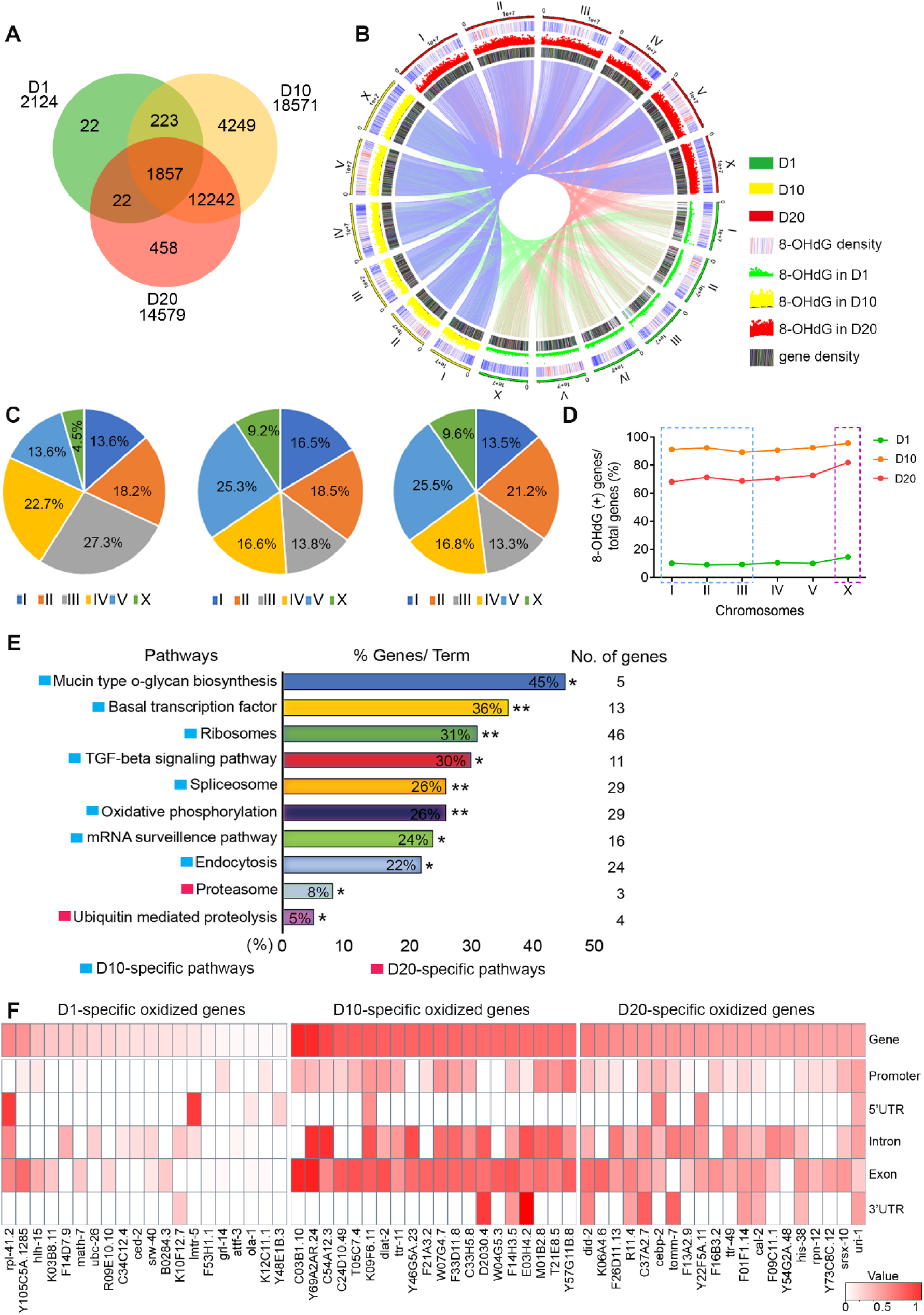
Oxidative modification patterns change with ageing. **A**. Overlap of genes with 8-OHdG modifications in *C.elegans* of different ages. **B**. The circos plots show the density of 8-OHdG modified genes. Gene density was quantified by the number of genes in every 100 kbp window. **C**. Distribution of age-specific 8-OHdG modified genes across different chromosomes. **D**. The dynamics of the chromosomal distribution of age-specific 8-OHdG modified genes among different age groups. Data shown are the % of 8-OHdG modified genes relative to a total number of genes. **E**. The functional enrichment of age-specific oxidized genes. Selected GO term enrichment for the genes exclusively oxidized in 10-day- and 20-day-old *C. elegans*. \**P*<0.05, \*\**P*<0.01; two-sided. **F**. The heatmap shows the genomic distribution of 8-OHdG sites in top-20 age-specific oxidized genes.

To investigate the functional intuitions of 8-OHdG modification in physiological ageing (Collado et al.; Lopez-Otin et al., 2013), we categorized age-specific 8-OHdG-modified genes using the *C. elegans* gene ontology (GO) resource (Ashburner et al., 2000; Van Auken et al., 2009). We noted that these genes were highly enriched in biosynthetic and transcriptional regulation pathways; in particular, D10-specific 8-OHdG-modified genes were enriched in bio-synthesis pathways including ribosomes, spliceosome, oxidative phosphorylation, endocytosis, basal transcription factor, TGF-beta signalling pathway, and mucin-type O-glycan biosynthesis, while D20-specific genes were enriched in proteolysis pathways (Fig. 3E). Interestingly, we noticed that top 8-OHdG modified genes identified in our samples were not yet functionally characterized so far (Table S4-6). These data highlight the importance of whole-genome 8-OHdG localization, which can facilitate to identify novel genes involved in redox mediated ageing. Upon analyzing the functional regions, we observed that 8-OHdG modifications are mainly present at the promoter and exon regions of 10- and 20-day-old worms, but, predominantly distributed at the intron regions of 1-day-old worms (Fig. 3F, Fig. S6D-F and Table S4-6); suggesting that 8-OHdG modification may affect gene regulation in an age-specific manner.

### The differentially 8-OHdG modified genes are annotated to longevity regulating pathways

Further, we examined the differentially oxidized genes (DOGs) between young and older worms (Fig. 4A). DOGs across different chromosomes demonstrated a similar trend in all groups with a comparatively high frequency in the sex chromosome (Fig. S7A). The differential 8-OHdG sites were highly observed in the promoter and exon regions (Fig. S7B-D). Gene Ontology analysis indicated that the sets of DOGs were enriched in multiple signalling pathways related to cellular processes, metabolism, and environmental information processing (Fig. 4B and Fig. S8A-C). Functional classification using KEGG databases indicated that DOG clusters were highly enriched in oxidative-stress related pathways, including FoxO, MAPK, ErbB, AGE-RAGE signalling, and longevity regulating pathways (Fig. 4C-F), suggesting that 8-OHdG modification indeed plays a key role in longevity regulation.

**Figure 4.**
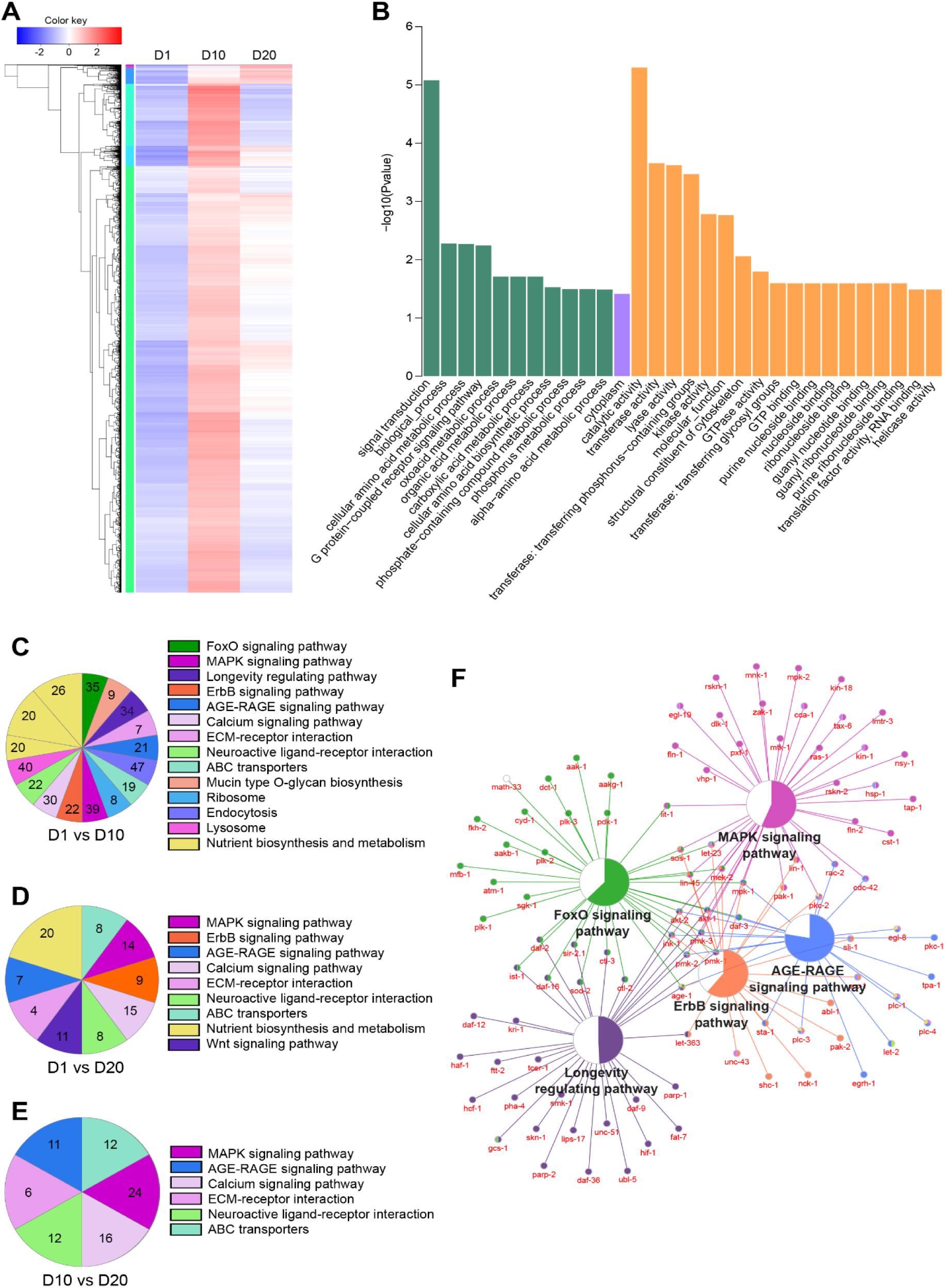
The differentially 8-OHdG modified genes are annotated to longevity regulating pathways. **A**. Heatmap of the 8-OHdG level of differentially oxidized regions (DORs) among different samples. The color key from blue to red indicates the relative 8-OHdG modification level in *k*-value from low to high, respectively. **B**. Major Gene Ontology (GO) category annotations of genes associated with 8-OHdG modification. Gene ontology analysis of differentially oxidized genes (DOGs, fold change modification bases in each gene between D1 *vs.* D10). **C-E**. Distribution of DOG clusters in different age groups. **C** indicates differentially oxidized pathways between D10 *vs.* D1; **D** indicates the comparisons between D20 *vs.* D1; **E** indicates D20 *vs.* D10. Portions of pie charts represent the proportion of genes in the pathway that are oxidatively modified within each biological process, using a *p*-value cutoff of 0.01. The number indicates the number of 8-OHdG modified genes in each pathway. **F**. The figure represents the top-5 signaling pathways that annotated DOGs. The Color portion of pie chart in each signaling pathway represents the percentage of genes in the pathway that are associated with 8-OHdG modification.

### The oxidative DNA damage in promoter regions is negatively correlated with the expression of pro-longevity genes

To identify the modification status of longevity regulating genes, we analyzed the 8-OHdG distribution in previously identified longevity regulating genes deposited at the animal aging and longevity database (AnAge) (Uno and Nishida, 2016). To examine the influence of oxidative modification on ageing, we divided longevity regulating genes into pro- and anti-longevity genes and plotted them against the occupancy of 8-OHdG in different age groups. Gene set enrichment analysis showed that the oxidative modification in pro-longevity and telomere maintenance genes increased with age (Fig. 5A and B). In young worms, we identified very few 8-OHdG positive sites in longevity regulating genes, however in the older worms, we found higher frequency of oxidative damage sites especially in pro-longevity genes, with predominantly localized in promoter regions (Table S7 and Fig. 5C). The 8-OHdG enrichment in longevity regulating genes among different age groups was further confirmed by 8-OHdG-IP-qPCR analysis (Fig. S9). Studies have suggested that age-associated changes in gene expression could be one of the key mechanisms of ageing (de Magalhaes et al., 2009; Harries et al., 2011; Kawahara et al., 2009); hence, to detect the relationship between 8-OHdG modification and expression levels of longevity regulating genes, we performed RNA-seq on different age *C. elegans*. As expected, the genome-wide transcriptional map analyzed from RNA-Seq data indicated an age-specific pattern of transcript expression in the *C. elegans* genome evidenced by differential expression profiles among different age groups (Fig. S10A-D). Similarly, the transcriptional profile of genes involved in top-5 signaling pathways enriched by DOGs also showed significant differences among diverse age groups (Fig. S11A and B). Further, correlation analysis showed a negative correlation between 8-OHdG modification in promoter with the expression of pro-longevity genes (Fig. 5D-F and Fig. S12A and B), suggesting that 8-OHdG modification at promoter regions may possess epigenetic potential.

**Figure 5.**
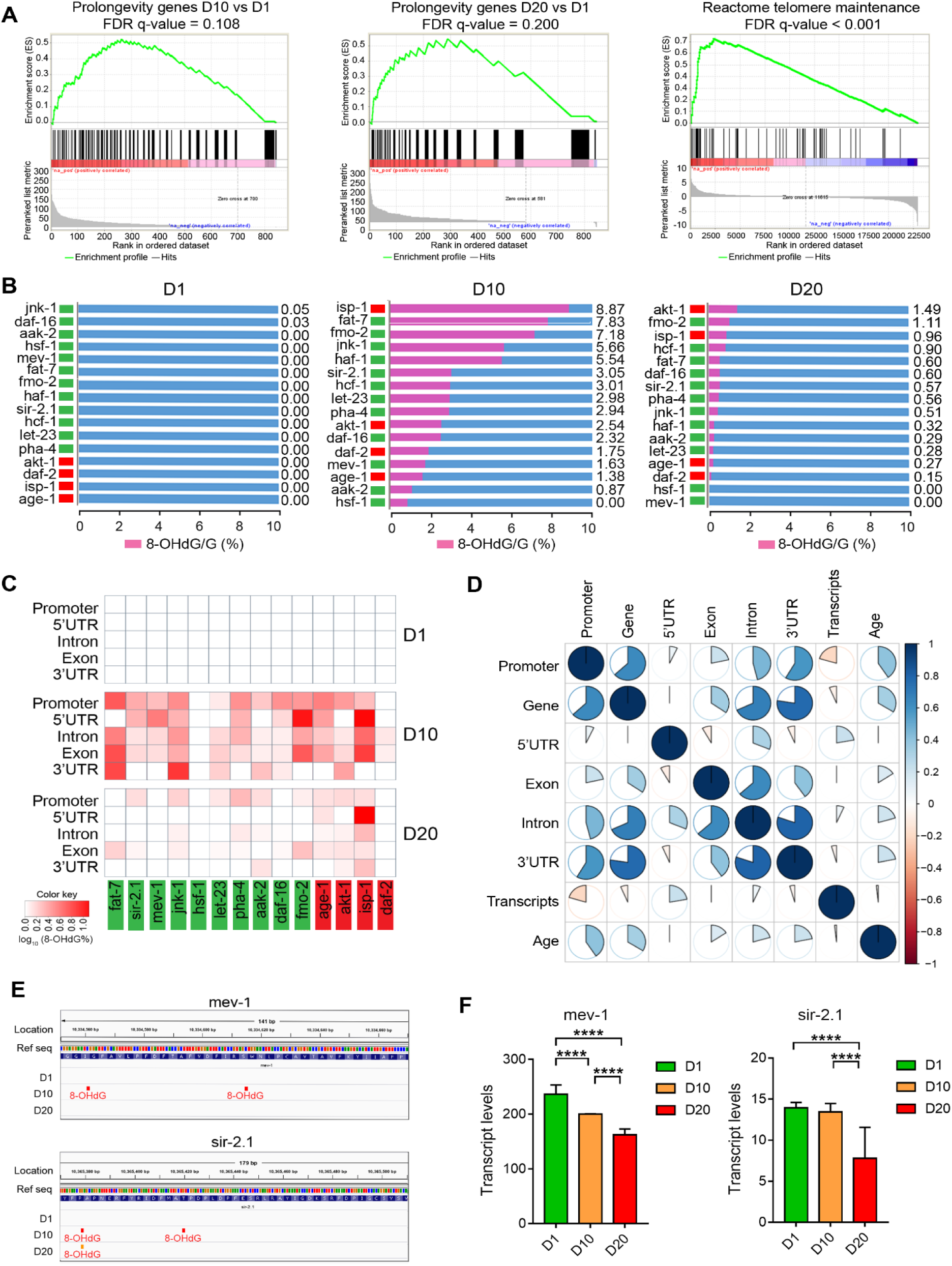
The oxidative DNA damage in promoter regions is negatively correlated with the expression of pro-longevity genes. **A**. The pro-longevity gene set enriched by the differential 8-OHdG modification between 10-day- *vs.* 1-day-old, and 20-day- *vs.* 1-day-old worms and telomere maintenance gene sets enriched by the differential 8-OHdG modification between 20-day- *vs.* 1-day-old worms. **B**. Comparisons of 8-OHdG modification levels in key ageing regulatory genes among different samples. Green indicates pro-longevity; red indicates anti-longevity. The genes are ordered by 8-OHdG modification levels. **C**. Representative figure showing the distribution of 8-OHdG modifications in functional regions of key ageing regulatory-genes. **D**. The correlation analysis for 8-OHdG modification and transcript expression of pro-longevity genes. The color scheme indicates the *Pearson* correlation coefficient, indicating the strength of the correlation between two variables. **E**. Genome browser view of representative loci in mev-1 and sir-2.1 genes across different samples. 8-OHdG sites are indicated color bars. **F**. The transcript expression levels of mev-1 and sir-2.1. The transcript level for each gene was estimated by averaging the fragments per kb exon model per million mapped reads (FPKM).

## Discussion

The underlying cause of ageing remains one of the central mysteries of biology. The oxidative stress theory of ageing convinces that both rates of ageing and life expectancy of an organism are largely influenced by the levels of accumulation of oxidative damage (Lin and Flint Beal, 2003). Hence, locating 8-OHdG at whole genomic level will provide a better understanding of the effects it has on normal ageing. *C. elegans* is one of the well-established premier model systems for aging research due to its short lifespan, relatively small genome size and simple body plan. In the current study, we mapped the whole-genome landscape of oxidative modification in *C. elegans* at different age periods and observed an agewise damage-driven reprogramming of pro-longevity genes. Our data set the stage for further investigation that will improve the understanding of the oxigenetic basis of ageing.

Our data showed that SMRT sequencing in Sequel platform is an efficient approach for mapping 8-OHdG sites at single-base resolution. As compared to other 8-OHdG sequencing methods, SMRT allows to assign the 8-OHdG mediated IPD signatures and motifs, which is different from other type of base modification under the high sequencing depth (Ardui et al., 2018). Besides, previous methods using second-generation sequencing require an ideal control sample, in which all the identified modification regions were based on mapping peaks relative to control. In addition, the absence of chemical tagging could improve the sequencing efficiency and accuracy, and maintain the physical and chemical integrity of DNA, providing more genuine information (Bryan et al., 2014; Clark et al., 2011; Ding et al., 2017; Poetsch et al., 2018; Schibel et al., 2010; Wu et al., 2018). Also, it can directly localize the medication and can identify individual bases and their modification simultaneously. Furthermore, SMRT sequencing can achieve higher accuracy for GC rich regions, where 8-OHdG modification frequently takes place (Wu et al., 2018). However, the drawback of this approach is the kinetic variation due to 8-OHdG modification or other unknown modifications that can influence the IPD values of both the upstream and downstream bases, increasing the false positive rate (Clark et al., 2011; Flusberg et al., 2010). Though we have filtered out the known 6mA and 4mC modification signals in our analysis, there are still many other known and unknown DNA modifications to be ruled out. Hence, further research addressing the accuracy of detection using integrated experimental and computational approaches is still required to efficiently map oxidative DNA damage at single-base resolution level.

The whole-genome 8-oxoG map in yeast suggests that greater oxidative damage distribution is determined by chromatin architecture, transcription activity, and chemical oxidation potential. In yeast genome, higher frequency of 8-oxoG was observed within heterochromatin and lower frequency in euchromatins, and in promoter and exon regions (Wu et al., 2018); whereas in mouse genome, higher accumulation of damage distribution occupied promoter regions (Ding et al., 2017). In contrast, oxidative damage distribution in human HepG2 cell line at the 10-300 kp resolution showed a reduced damage in functional regions such as coding, promoters, and transcription factor binding sites (Poetsch et al., 2018). In contrast, our finding at single-base resolution indicated that oxidative modification is highly distributed in non-functional regions in young worms, but for older worms, oxidative damage sites are more frequently identified at functional regions. This discrepancy likely results from age-specific- or regioselective distribution in oxidative damage sites and repair capacity, due to species specificity or due to the influence of environment and life-history which shapes the oxidative damage distribution through the evolutionary aspect (Speakman et al., 2015). Another factor to consider is that all previous studies observed the oxidative DNA damage at a single age period or a designated time point; and since oxidative DNA damage itself is a dynamic process, the changes in oxidative modification pattern throughout the life span might have been missed. Therefore, further studies conducted at different age periods, with larger samples size, and strict environmental controls are required to draw the representative landscape of whole-genome oxidative DNA modification across different species.

Although several experimental findings have supported the concept that ageing-associated cellular changes are due to the accumulation of oxidative DNA damage (Bergamini et al., 2004; Chen et al., 2007; De Bont and van Larebeke, 2004; Lenaz, 1998; Lin and Flint Beal, 2003), many recent studies suggest that it could not be the sole cause (Liochev, 2013; Pérez et al., 2009; Sanchez-Flores et al., 2017) and highlighted the complexity of the mechanisms behind ageing. Thus, an integrated epidemiological, genetics and epigenetics approach is required for the betterment of biological ageing research (Fallin and Matteini, 2009; Speakman et al., 2015). Previous studies paid much attention on the impact of oxidative DNA damage on ageing associated disease including Alzheimer, cardiovascular diseases and carcinogenesis and undermined the role of oxidative DNA damage in physiological ageing. Interestingly, our data shows that the frequency of genomic oxidative DNA modification is the highest in middle age worms and slowly declines as they age, suggesting that that higher oxidative modification sites are not proportionally correlated with the rate of ageing. This could be partially explained by age-driven changes in DNA damage tolerance along with their response to oxidative stress (Pilzecker et al., 2019). Although middle aged worms experienced higher levels of oxidative stress, they might have better damage tolerance, which can ameliorate the DNA damage-related biological consequences and can achieve the normal equilibrium at a faster pace relative to the older worms (Hekimi et al., 2001). In addition, one recent study has shown that transient increase in naturally occurring reactive oxygen species during early development increases stress resistance and improves redox homeostasis and consequently brings a positive effect on lifespan regulation (Bazopoulou et al., 2019). However, more studies, including mechanistic ones, are needed for the deeper understanding of the impact of oxidative DNA damage to the physiological ageing and health spans.

The differential 8-OHdG modified genes are highly enriched in age regulating pathways, including longevity regulating pathway, and FoxO, ErbB, MAPK, and AGE-RAGE signaling pathways. These pathways are consistent with the previously reported ageing-dependent signaling pathways (He et al., 2014; Lapierre and Hansen, 2012), indicating the significance of oxidative modification in ageing processes. The rate of 8-OHdG modification in key ageing regulating genes is different among diverse age groups. These modifications are mainly observed in exon and promoter regions. Studies have reported that age-associated changes in gene expression are one of the key underlying mechanisms behind the physiologic consequences of ageing (de Magalhaes et al., 2009; Harries et al., 2011); favourable to these reports, we observed an age-specific pattern of transcript expression in the *C. elegans* genome. The transcriptional profile of genes involved in pathways enriched by DOGs showed significant differences among different age groups. These findings are highly consistent with previous studies where 8-oxoG accumulates in the promoters of regulatory genes such as VEGF (Pastukh et al., 2015) and p53 (Hyun and Jang, 2014; Ou and Schumacher, 2018) upon oxidative response and consequently regulate gene expression. Studies demonstrated that oxidative DNA damage in the promoter region could halt transcription and replication (Cogoi et al., 2018; Tornaletti et al., 2004), resulting in cellular dysfunction that promotes cellular senescence and apoptosis (Ou and Schumacher, 2018). Consistently, our data unveiled that 8-OHdG modification in promoter regions was negatively correlated with the expression of pro-longevity genes, indicating that 8-OHdG at promoter regions may have epigenetic potential (Fleming et al., 2017). In young worms, we identified almost none oxidative damage sites in longevity regulating genes, however, in the older worms, we found higher frequency of oxidative damage sites especially in pro-longevity genes, suggesting that oxidative modification in pro-longevity genes may have a robust effect on lifespan determination as compared to anti-longevity genes. Furthermore, our data specify that age-associated changes in oxidative DNA-damage driven transcript repression exclusively occurs during adulthood.

In summary, our results delineate the feasibility of measuring oxidative sites using PacBio’s Sequel platform. Our findings reveal that oxidative DNA damage can impact longevity regulation in an age-specific manner. Accumulation of oxidative DNA damage in pro-longevity genes of older worms can repress their expression, mediating organismal ageing. This concept highlights the previously unappreciated, epigenetic role of oxidative DNA modification on longevity regulation. However, further research addressing the accuracy of 8-OHdG detection using an integrated experimental and computational approach is still required to improve the mapping of oxidative DNA damage at the single-nucleotide resolution level. We believe that our findings provide a comprehensive data on the oxidative DNA modification map and set the stage to improve our understanding of the molecular mechanisms of 8-OHdG in ageing, thereby providing the basis to prolong the youthfulness and lifespan.

## Materials and Methods

### Synthetic 8-OHdG containing oligonucleotide sequence

Custom oligonucleotides containing 8-OHdG bases were purchased from Takara Bio Inc. The sequence information is 5’ -GTCGTACGACGTTATTCGTTTCGTCCGACGATC(x)T ACGACGTTATTCGTTTCGTCC(x)ACGACGGGCTCGGAACGAAAGTTCCGAGCCCGTCGTCGGACGAAACGAATAACGTCGTACGATCGCGGACGAAACGAATAACGTCGTACGACT-3’ for the forward strand and 5’-AGTCGTACGACGTTATTCGTTTCGTCCGACGATCGTACGACGTTATTCGTTTCGTCCGACGACGGGCTCGGAACTTTCGTTCCGAGCCCGTCGTCGGACGAAACGAATAACGTCGTACGATCGTCGGACGAAACGAATAACGTCGTACGAC-3’ for the reversed strand. The “x” denotes for 8-hydroxy-2’ -deoxyguanosine (8-OHdG).

### Preparation of plasmid DNA

The Haemagglutinin (HA)-tagged pcDNA3.1(+) plasmid DNA was amplified in Escherichia coli DH5α and extracted with EasyPure HiPure Plasmid MaxiPrep Kit (TransGen). The linearized plasmid used for hydrogen peroxide treatment and library construction was cut by EcoRV (Takara) to generate blunt-ends. To prevent 8-OHdG formation during the procedure, we added 1% of 100 mM deferoxamine to the reaction buffer. For DNA purification, 1/10 volume of 5 M NaCl and 5x volume of 100% ethanol were added and samples were placed at −20°C for at least 30min. Samples were spun at 12,000 rpm at 4°C for 30 minutes. The supernatant was removed. Equivoluminal cold fresh 75% ethanol was added and samples were again spun at 13,000 rpm for 10 minutes. The supernatant was discarded, and the pellet was allowed to dry before being resuspended in DNase-free water.

### Plasmid DNA oxidation by hydrogen peroxide treatment

Fenton’s reaction was used to induce oxidative damage (Wang et al., 2015). Final concentrations of 10 μM H_2_O_2_ and 2.5 μM Fe^2+^ were mixed with 70 μg/ml of linearized pcDNA3.1-HA plasmid diluted in nuclease-free water. The Fenton’s reaction was incubated for 1 hour at 37°C. The reaction can be blocked by placing the samples on ice or purifying the DNA immediately, and the purification method was as described above.

### Base excision reaction by Ogg1 treatment

The standard assay of mOgg1 reaction was prepared as described (Zharkov et al., 2000) with slight modification. Briefly, reaction mixtures included 10ug of DNA, 25 mM sodium phosphate, pH 7.5, 100 mM NaCl, 2 mM Na-EDTA, and 10 ug mOgg1 in a total volume of 1 ml. Reactions were initiated by adding enzyme and allowed to proceed for 30 min at 37°C.

### Plasmid DNA point mutation

A point mutation was constructed with the Fast mutagenesis system kit (TransGen) and performed as the manufacturer’s described by designing divergent primers.

MUT-random-forward: 5’-CAGTTGGGTGCACGACTGGGTTACATCGAAC-3’;

MUT-random-reverse: 5’-GTCGTGCACCC AACTGATCTTCAGCATCTTTTAC-3’;

MUT-2427-forward: 5’-CATGTAACTCGCCTTCATCGTT GGGAACCGG-3’;

MUT-2427-reverse: 5’-GAAGGCGAGTTACATGATCCCCCATGTTGTG-3’.

### Worms culture and synchronization

The worms were cultured as others described with slightly modifications (Stiernagle, 2006). E. coli OP50 bacteria were cultured in Luria broth (LB) overnight at 37°C on 180rpm. The overnight cultured OP50 (100 μl) was seeded on nematode growth media (NGM, Luria broth, Agarose powder) plates and incubated overnight at 37°C for another 6 hours. The prepared NGM plates were used for wild-type strain N2 maintaining at 20°C. For *C. elegans* synchronization, worms were washed with sterile H_2_O and collected in a centrifuge tube, A total of 3.5 ml of sterile H_2_O, and then 1.5 ml of NaOH and bleach mix (0.5 ml 5 N NaOH,1 ml bleach) was added to the tube. The tube was vortexed for 20 seconds every 2 minutes for a total of 10 minutes and then centrifuged for 30 seconds at 1300g to pellet released eggs. The eggs were resuspended with H_2_O and transferred in a new prepared NGM plate.

### *C. elegans* sample preparation

The hatched larvae were transferred to a newly prepared NGM plate and harvested the 1-day adult worms for gDNA extraction. To prevent the progeny from hatching, 5-fluoro-2’-deoxyuridine (Fudr) was added to the plates at a final concentration of 50 µM (Heehwa G. Son, 2017). 1-day adult worms were placed on Fudr plates, and then transferred to Fudr plates every 3-4 days. After 10- and 20-days, worms were collected for gDNA extraction (Fig. S4), respectively. We collected a total of 30 plates for 1-day old, 30 plates for 10-day old and 60 plates for 20-day-old groups. We further subdivided the collected worms into two tubes for DNA extraction and RNA extraction, respectively in order to maintain the biology and batch consistency of *C. elegans* between DNA and RNA sequencing samples.

### Worm gDNA extraction

We extracted total DNA of the *C.elegans* with the FastPure Cell/Tissue DNA Isolation Mini Kit (Vazyme) according to the manufacturer’s instructions. To prevent 8-OHdG formation during the procedure, we added 1% of 100 mM deferoxamine to the reaction buffer. The concentration and quality of the resulting DNA were checked using the Nanodrop and Qubit dsDNA high sensitivity assay kit (Invitrogen).

### Library preparation and SMRT sequencing

The genome sequencing was performed using Pacbio SMRT sequencing protocol as described (Xincong Kang, 2017) with modifications. First, the integrity, quality, and concentration of total DNA were analyzed by agarose gel electrophoresis, NanoDrop spectrophotometer (Thermo Fisher Scientific), and Qubit 3.0 fluorometer (Thermo Fisher Scientific). To prevent 8-OHdG formation during the library construction, we added 1% of 100 mM deferoxamine to the extracted DNA. DNA was haphazardly sheared to fragments with an average size of 5 kb (for plasmid gDNA) and 20 kb (for *C. elegans* gDNA) by using g-TUBE. Sheared DNA was then DNA end repaired. SMRTbell templates were obtained by ligating the blunt hairpin adapters to the ends of the repaired fragments, omitting repair DNA Damage step to avoid the potential loss of oxidative modification mediated by DNA repair mix, followed by the addition of exonuclease to remove failed ligation products. After annealing the sequencing primer and binding polymerase to SMRTbell templates, a total of 5 (1 each for 0 µM, 10µM, and 10µM+Ogg1 treated and two point-mutated samples) SMRT cells were sequenced on a PacBio SMRT sequencing platform. For *C. elegans* DNA, it was mechanically sheared by a Covaris g-TUBE device at 4,800 rpm for 2 minutes. SMRTbell templates were bound to v3 primers and polymerases using Sequel^TM^ Binding Kit 2.1. Polymerase bound complexes were purified with SMRTbell Clean Up Column Kit and sequenced by diffusion loading method. We sequenced the plasmid (using 8 SMRT cells) and *C. elegans* gDNA libraries (using 13 SMRT cells: 2 for D1; 8 for D10; 3 for D20) in a PacBio Sequel platform (Instrument ID: 54188) with a movie length of 600 minutes at the Center for Molecular Genetics, Institute for Translational Medicine, Qingdao University (Qingdao, China).

### RNA-Seq

RNA-seq was performed as described (Wang et al., 2009) with slight modifications. Briefly, 1-day-old (n=30 plates), 10-day-old (n=30 plates) and 20-day-old (n=60 plates) wild-type *C. elegans* were harvested for total RNA extraction (Fig. S4) using the Trizol reagent (Invitorgen). The amount and quality of RNA were tested by Qubit RNA Assay Kit (Life Technologies), gel electrophoresis and Agilent Bioanalyzer 2100 system. NEBNext Ultra Directional RNA Library Prep Kit for Illumina (NEB) was used to construct the library according to the manufacturer’s instructions, the sequencing libraries were prepared. Poly-A-containing mRNA was isolated from the total RNA by poly-T oligo-attached magnetic beads, and then fragmented by RNA frag-mentation kit. The cDNA was synthesized using random primers through reverse transcription. After the ligation with adaptor, the cDNA was amplified by 15 cycles of PCR, and then 200-bp fragments were isolated using gel electrophoresis. We prepared 2 libraries for each age group, and each library was sequenced in 2 separate lanes by an Illumina NovaSeq instrument in Shanghai Majorbio Biopharm Technology Co. Ltd. (Shanghai, China).

### HPLC and HPLC-MS/MS

8-oxo-7,8-dihydroguanosine (8-OHdG) was analyzed by HPLC and HPLC-MS/MS as previously described (Allan Weimann, 2002; Henriksen et al., 2009; Wang et al., 2015) with modifications. In brief, 10μg gDNAs were completely dissolved in 8.5 μl of 300 μM deferoxamine mesylate (DFOM). Addition of 3.2 μl Nuclease P1 (stock of 1.25 U/μl in 300 mM sodium acetate, 0.2 mM ZnCl2, pH 5.3, frozen at -20°C) was followed by 13.7μl alkaline phosphatase (0.365U/μl), and hydrolysis took place at 50°C for 60 min. After centrifugation for 10 s (2000g), the hydrolysate (100 μl) was transferred to a Micropure-EZ filter (Millipore) and centrifuged at 14,000g for 60 s for enzyme removal. The hydrolysate was placed into an autosampler vial for HPLC-MS/MS analysis using Agilent Technologies 1290 Infinity II. The samples were separated on a C18 reverse phase column (Agilent Poroshell 120 EC-C18 27μm). The mobile phase composition was ammonium acetate (20 mmol/L, pH 6.5)-methanol (90:10) at a flow-rate of 0.4 ml/min. The 8-OHdG standard was from Sigma-Aldrich. Mass spectrometric detection was performed on LC-ESI-MS system (Agilent Technologies 6460 Triple Quad LC/MS) equipped with an ESI ion source operated in the positive mode. The MS setting parameters were: Gas Temp 350℃, Gas Flow 10 l/min, Nebulizer 35psi, Capillary Positive 4000V, Capillary Negative 3500V, Frag=175V. The instruments were controlled by the Agilent MassHunter Workstation Software. The MS showed a Counts vs. Mass-to-Charge m/z for 8-OHdG.

### 8-OHdG-IP-qPCR

The worm DNA sample was sonicated to produce ∼250bp fragments. The worms’ DNA was incubated with specific anti-8-Hydroxy-2’-deoxyguanosine antibody (Abcam) in immunoprecipitation buffer (2.7 mM KCl, 2 mM potassium phosphate, 137 mM NaCl, 10 mM sodium phosphate, 0.05% Triton X-100, pH 7.4) for 2h at 4°C. The mixture was then immunoprecipitated by Protein A/G Plus-Agarose (Santa Cruz) that was pre-bound with bovine serum albumin (BSA) at 4°C for 2h. After extensive washing, the bound DNA was eluted from the beads in elution buffer (50 mM Tris, pH 8.0, 1% SDS, 10 mM EDTA) at 65°C, treated with proteinase K and purified using QIAquick PCR Purification Kit (Qiagen). Fold-enrichment of each fragment was determined by quantitative real-time PCR. The primers were listed in Supplementary Table S8.

### Dot blot analysis

Dot blot assay was performed as others described with modifications (Greer et al., 2015b). 2500ng DNA samples were loaded per dot on nylon membranes, λDNA was loaded as negative control. Membranes were allowed to air dry and then DNA was autocrosslinked in a UV stratalinker (80J) for 2 times. The membrane was then blocked for 2 hours in 3% BSA at room temperature. Membranes were probed for 1 hour at room temperature and overnight at 4°C with primary antibody (Anti-8-Hydroxy-2’-deoxyguanosine antibody, 1:1000, Abcam) in 3% BSA. Blots were washed 3 times for 15 minutes with TBST then probed with secondary antibody (Goat anti mouse IGg(H+L) HRP, 1:1000, TransGen) in 3% BSA for 1 hour at room temperature. Blots were washed 3 times for 15 minutes with TBST and ECL was applied and film was developed.

## Quantification and statistical analysis

### 1. Mapping and characterizing 8-OHdG modifications

#### (1) Genome references

SMRT-seq data were mapped to the appropriate genomes using BLASR via SMRTportal. The *C. elegans* data were mapped to C. elegans genome version *WBcel235* (ftp://ftp.ensemblgenomes.org/pub/metazoa/release-44/fasta/caenorhabditis_elegans).

#### (2) Pre-processing of SMRT sequencing data for IPD analysis

We followed the pre-processing steps as implemented in SMRTportal (Greer et al., 2015a). In brief, an initial filtering step removes all subreads with ambiguous alignments (MapQV<240), low accuracy (<75%) or short-aligned length (<50 bases). Next, an additional filtering step removes the subread IPD values from the mismatched positions with respect to the reference sequence. To prevent any potential slowing of polymerase kinetics over the course of an entire read, subread IPD normalization is done by dividing subread IPD values by their mean.

#### (3) Identification of 8-OHdG modification

Each of the raw data in h5 format were first aligned to reference genome using pbalign in base modification identification mode. Then the post-aligned datasets were merged and sorted by using pbmerge and cmp5tool. The depth of sequencing dataset of all samples is above 100x coverage (Table S1), higher than the recommended coverage for detecting 8-OHdG modification in SMRT portal. Finally, the 8-OHdG was identified using ipdSummary.py script according to published protocol with slight modification (Clark et al., 2011; Flusberg et al., 2010). In brief, the single-molecule base modification analysis used to query the underlying heterogeneity of modifications across individual molecules is analogous to the standard base modification analysis supported in SMRT Analysis. The IPD ratio per given position is calculated by averaging the IPD values across all the subreads from multiple molecules before comparing with the in-silico control IPD. The statistically significant between IPD value at each position with the in-silico control is determined by the modification score (−log_10_ *p*-value). Reads were filtered for having five or more subreads and required three or more IPDs with the preceding and following base correction for computation of the single-molecule IPD ratio. We then further filtered 8-OHdG sites with more than 25x coverage and <0.05 FDR as described above using customized scripts in Python. For motif identification, we then extracted 2 or 3bp from the upstream and downstream sequences and consensus sequence were analyzed for the identification of potential motif using customized Perl script. Motif analysis on consensus sequences next to 8-OHdG modification in *C. elegans* genome was performed using the Motif Elicitation by Expectation Maximization (MEME) algorithm (Bailey et al., 2009). Circos plot, gene annotations, genomic distribution and genomic density of 8-OHdG sites were analyzed by customized scripts in R, Python and Perl with the assistance of Hyde Resource Biotechnology, Inc. Qingdao.

#### (4) Estimation of false discovery rate (FDR) for single nucleotide-level 8-OHdG calls

Considering each molecule separately, the IPD values (post-filtering) are grouped by their strand and mapped genomic position, and the mean value is calculated. At each genomic position of a single strand, the mean IPD values for each molecule follow the Gaussian distribution based on Central Limit Theory. The FDR corresponding to a specific threshold on a given statistical measure (e.g. IPD ratio, *t*-test *p-*value or identification Qv) is estimated by comparing the global distribution of the measure obtained from the native DNA sample with that from the G sites across the genome. Specifically, the FDR is calculated for each motif, FDRs can be estimated for single nucleotide-level 8-OHdG calls only among the G sites corresponding to the motif across the genome. We generated the corresponding ROC curves by sliding a threshold value across the full range of observed IPDs. We computed the true positive rate for each threshold as the fraction of oxidized modification observations with an average IPD larger than the threshold. Similarly, we calculated the false positive rate by the fraction of non-oxidized observations with an average IPD larger than the threshold. Genome-wide distribution of 8-OHdG was mapped by R software circlize package.

#### (5) Genomic region annotations and calculation of 8-OHdG modification levels

We classified the genomic regions into 5 classes including 5’UTR, promoters, exons, introns, 3’UTR and annotated them as previously published *C. elegans (WBcel235)* reference deposited in Ensemble Metazoa database (ftp://ftp.ensemblgenomes.org/pub/metazoa/release-44/fasta/caenorhabditis_elegans/dna/). For each annotated genomic region, the 8-OHdG level was calculated as the average 8-OHdG sites per total G bases covered within the region.

### 2. Differentially oxidized regions (DORs)

DORs were identified by comparing the number of 8-OHdG sites within the same genomic length and region between the two samples. The 8-OHdG modification sites in 100kbp windows are counted using in-house Perl scripts and we assigned these regions as DORs only if the 8-OHdG sites in one sample is greater than or less than another sample, with a significant *p*-value < 0.05 given by multiple Student’s t-test and a Benjamini–Hochberg false discovery rate (FDR) < 0.05.

### 3. Differentially oxidized genes (DOGs)

DOGs were identified by comparing the number of 8-OHdG sites observed in the same gene annotated between two samples. The 8-OHdG modification sites in the whole length of the encoded gene was counted using customized Perl scripts and we assigned these genes as DOGs only if the 8-OHdG sites in the gene region of one sample was greater than or less than that of another sample, with a significant *p*-value < 0.05 given by multiple Fisher exact test and a Benjamini–Hochberg false discovery rate (FDR) < 0.05; we assigned these genes as age-specifically oxidized gene only if the 8-OHdG sites were exclusively present in the annotated gene region of one sample compared to that of another sample.

### 4. Gene Ontology (GO) and KEGG (Kyoto Encyclopedia of Genes and Genomes) Analysis

GO analysis was performed using Goatools (https://github.com/tanghaibao/GOatools), the *p*-value was corrected by Bonferroni, Holm, Sidak methods false discovery rate) with a significant *p*-value cutoff of 0.01 and a Benjamini–Hochberg false discovery rate (FDR) < 0.05 (Gene Ontology, 2015). KEGG analysis was performed using KOBAS (http://kobas.cbi.pku.edu.cn/home.do) and cytoscape ClueGO package, with a significant *p*-value cutoff of 0.01 and a Benjamini–Hochberg false discovery rate (FDR) < 0.05 (Gene Ontology, 2015; Kanehisa and Goto, 2000).

### 5. RNA-Seq transcriptomic analysis

#### (1) Read mapping

The raw paired end reads were trimmed and quality controlled by SeqPrep (https://github.com/jstjohn/SeqPrep) and Sickle (https://github.com/najoshi/sickle) with default parameters. Sequences with primer concatemers, weak signal, and/or poly A/T tails were culled. Clean reads were aligned to the *C. elegans* (ftp://ftp.ensemblgenomes.org/pub/metazoa/release-43/fasta/caenorhabditis_elegans/dna/) reference using TopHat (http://tophat.cbcb.umd.edu/, version2.0.0) software (Ghosh and Chan, 2016; Trapnell et al., 2009). The mapping criteria of bowtie was as follows: sequencing reads should be uniquely matched to the genome allowing up to 2 mismatches, without insertions or deletions. Then the region of gene was expanded following depths of sites and the operon was obtained. In addition, the whole genome was split into multiple 15kbp windows that share 5kbp. New transcribed regions were defined as more than 2 consecutive windows without overlapped region of gene, where at least 2 reads mapped per window in the same orientation. The mapped reads were assembled by Cufflinks v2.1.1. Cuffdiff was used to calculate FPKMs of coding genes (Trapnell et al., 2012).

#### (2) Differential expression analysis and Functional enrichment

To identify DEGs (differential expression genes) between two different samples, the expression level of each transcript was calculated according to the fragments per kilobase of exon per million mapped reads (FRKM) method. RSEM (http://deweylab.biostat.wisc.edu/rsem/) was used to quantify gene abundances. R statistical package software EdgeR (Empirical analysis of Digital Gene Expression in R, http://www.bioconductor.org/packages/2.12/bioc/html/edgeR.html) was utilized for differential expression analysis. In addition, functional-enrichment analysis including GO and KEGG were performed to identify which DEGs were significantly enriched in GO terms and metabolic pathways at Bonferroni-corrected *p*-value ≤0.05 compared with the whole-transcriptome background. GO functional enrichment and KEGG pathway analysis were carried out by Goatools (https://github.com/tanghaibao/Goatools) and KOBAS (http://kobas.cbi.pku.edu.cn/home.do).

### 6. The integrated analysis of 8-OHdG modification in different genomic regions, gene expression and aging

We calculated the 8-OHdG sites in 5 different genomic regions as described above. The log2 of the gene expression levels (RPKM) of RefSeq genes were calculated as described (Trapnell et al., 2012). We plotted the Pearson correlation coefficients (r) between 8-OHdG levels of each genomic region/gene expression and different age group using R Bioconductor software packages (https://bioconductor.org/packages/3.9/bioc/).

### 7. Statistical Analysis

Data were analyzed by two-way ANOVA using Bonferroni posttest (Prism 5.0). All of the data are presented as the mean ± SEM and represent a minimum of three independent experiments.

## Acknowledgments

We thank to Hyde Resource Biotechnology, Inc. (Qingdao) for helpful advice on genomic sequencing and data analysis. DNA sequencing was performed at Qingdao University, Institute for Translational Medicine, Center for Molecular Genetics and data analysis was completed at Qingdao University, Institute for Translational Medicine, Center for Bioinformatics. This project was supported by the Major Research Program of the National Natural Science Foundation of China (Grant number 91849209), National Natural Science Foundation of China Research Fund for International Young Scientists (Grant number 81850410551), Natural Science Foundation of Shandong Province (Grant number ZR2019BH 014).

## Author’s contributions

L.H.H.A., and P.L., conceived and designed the study. L.H.H.A., Z.-Q.L., Z.L., Z.Y., X.C, and J.G., prepared the samples; L.H.H.A., P.S., and Z.Z. prepared the Ogg1 reaction assays; L.H.H.A. performed library construction and genomic sequencing. L.H.H.A., and P.L. analyzed the data; X.C., Z.-Q.L., and Z.L. helped to analyze the data. L.H.H.A. wrote the manuscript; L.H.H.A., P.L., and Y.W., edited and submitted the manuscript. All authors discussed and finalized the manuscript for submission.

## Supplementary Figures

**Figure S1.**
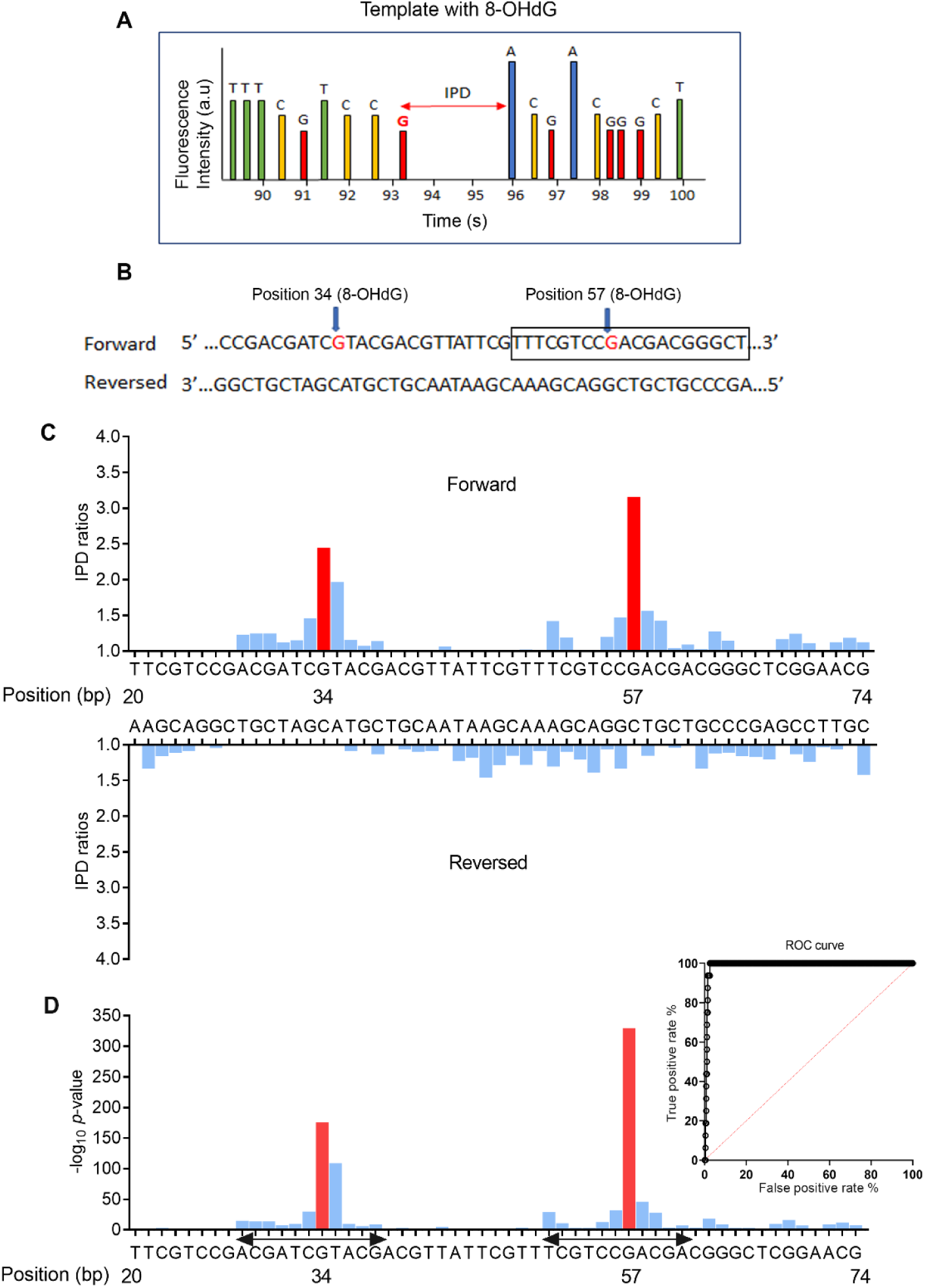
8-OHdG modification captured by SMRT sequencing. **A**. Schematic illustration of IPD prolongation in the presence of 8-OHdG modification. **B and C**. 8-OHdG modification captured by SMRT sequencing. To calibrate the method, we used an oligonucleotide containing two 8-OHdGs and sequenced in SMRT *RSII* platform. **B** shows the nucleotide sequence. **C and D** show the mean IPD ratio and modification score (−log_10_ *p*-value) at each position and putative 8-OHdG event detected at 8-OHdG sites. The IPD ratio of each base position was determined by comparing the ratio of the IPD at a site in the native sample to the IPD at a site in an in-silico control (Schadt et al., 2013) after measuring n=465 times in average. After filtering the sequencing depth and modification score using customized python scripts, putative 8-OHdG events were detected at the positions 34 and 57 of the synthetic oligonucleotide with slight IPD ratios changes next to the modification site. Receiver operating characteristic (ROC) curves, based on the IPD distributions from the positive oxidation signal parameterized by IPD threshold and medication score, for assigning oxidation status of guanosine nucleotide.

**Figure S2.**
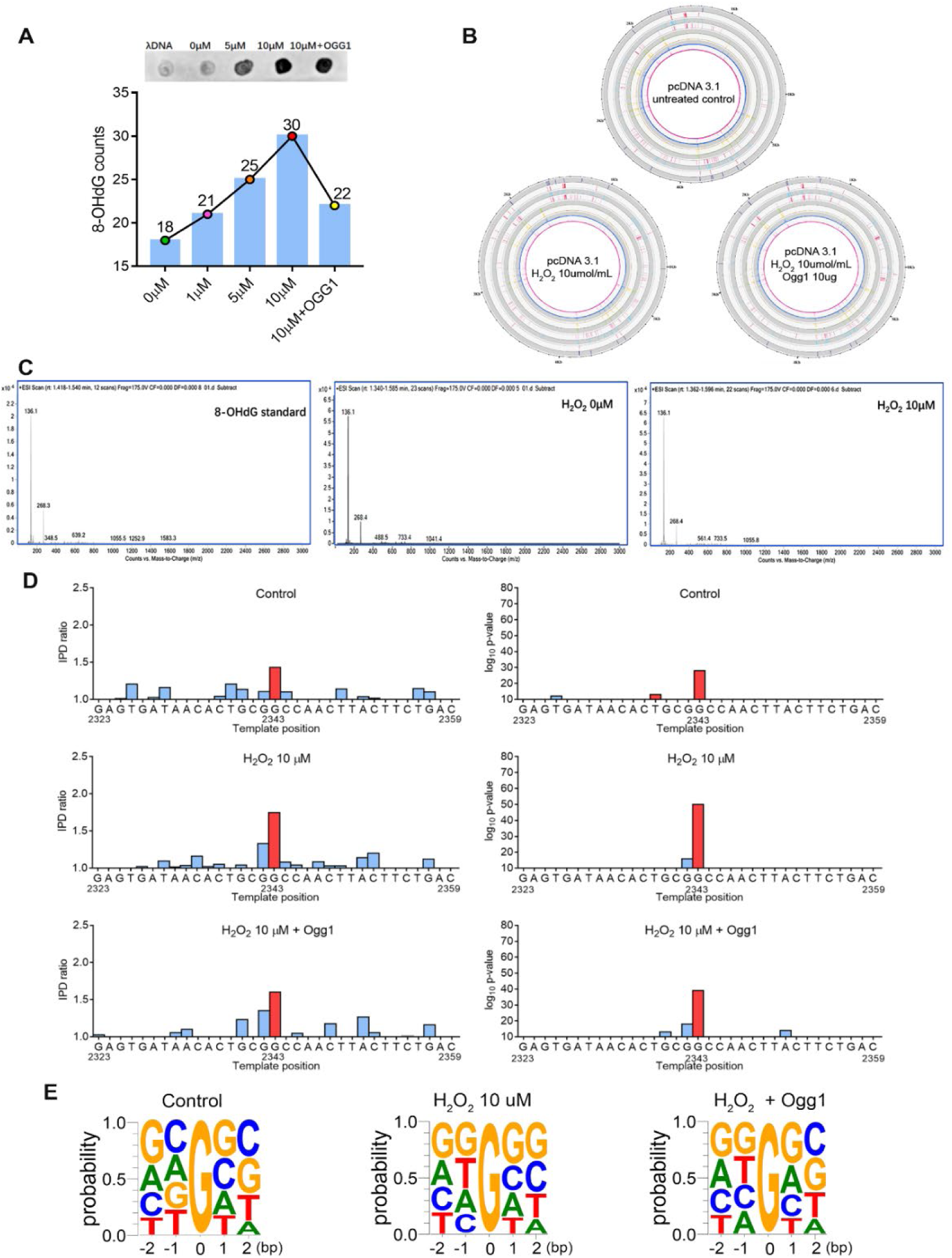
SMRT technology is feasible to detect 8-OHdG modification at whole-genome level in plasmid genome. **A.** The dot-blot analysis and 8-OHdG counts identified in different plasmid samples by SMRT sequencing. The plasmid genomic DNAs were treated with indicated dose of H_2_O_2_ and 10μg of Ogg1, and the presence of 8-OHdG was detected (n=2). **B**. Circos plots showing the distribution of 8-OHdG bases detected in different plasmid samples. From inner to outer circles: 1^st^ and 2^nd^ indicate IPD ratios, 3^th^ and 4^th^ represent the coverage of each position detected, 5^rd^ and 6^th^ indicate the modification score (−log_10_ *p*-value) for each position tested, and 7^th^and 8^th^ denote 8-OHdG modification for the forward and reversed strands, in untreated control, hydrogen peroxide (H_2_O_2_) treated and oxoguanine glycosylase (Ogg1) treated samples. **C.** The presence of 8-OHdG in each sample was confirmed by HPLC associated with tandem Mass Spectrometry (HPLC-MS/MS). D shows the mean IPD ratio and modification score (−log_10_ *p*-value) at each position of one 8-OHdG event detected in plasmid genome. **E**. Motif analysis based on consensus sequences (−2bp and +2bp) next to 8-OHdG sites detected under different conditions: untreated control, hydrogen peroxide (H_2_O_2_) treated and oxoguanine glycosylase (Ogg1) treated samples.

**Figure S3.**
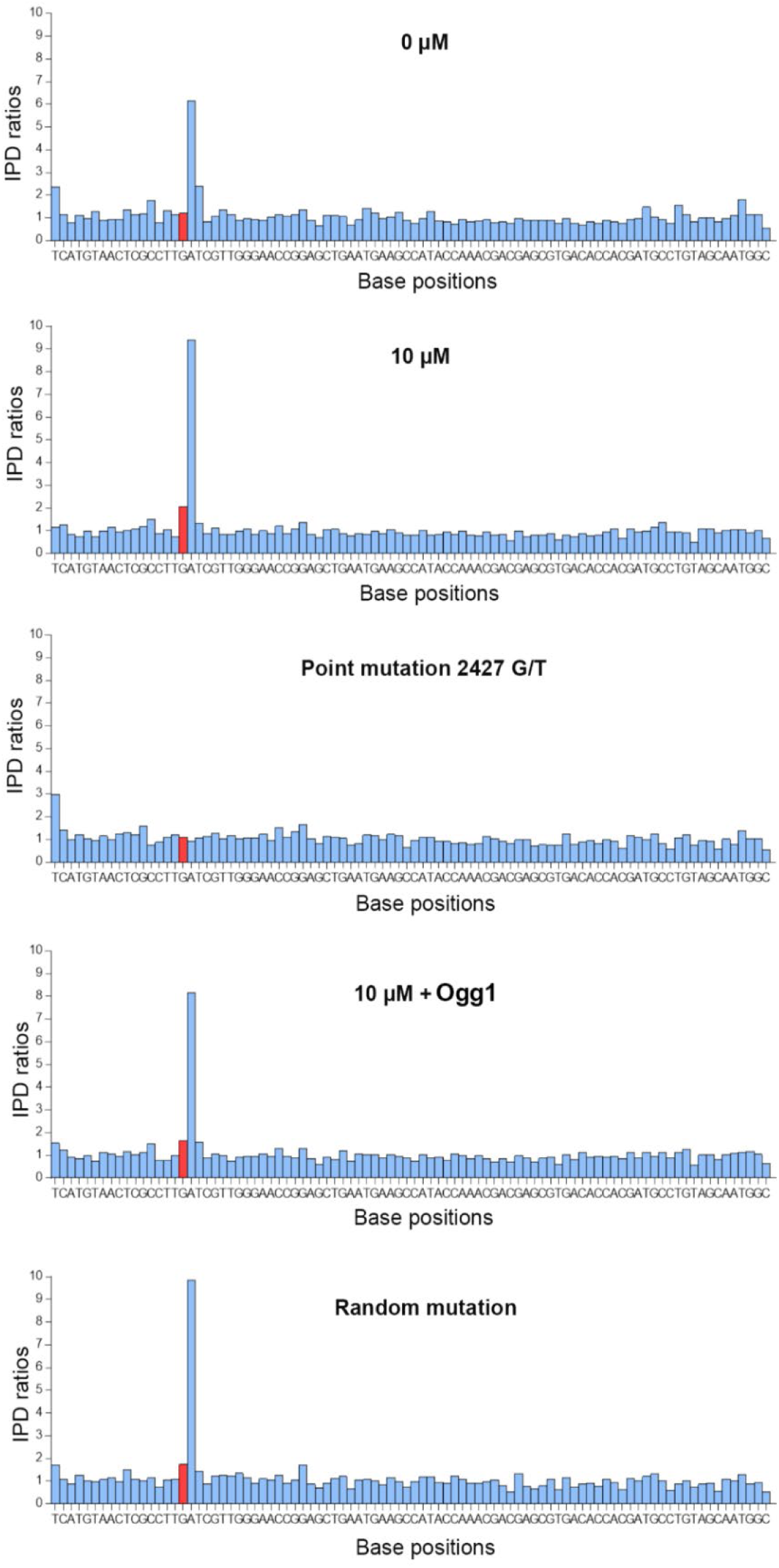
IPD ratios of 8-OHdG and its surrounding bases in the plasmid genome. Representative IPD ratio plots show one instance of 8-OHdG modification detected in WT, H_2_O_2_-treated, and H_2_O_2_-treated site-directed 2427 G/T mutation, Ogg1 and random mutation samples. No statistically significant modification signal was detected when G residue was mutated.

**Figure S4.**
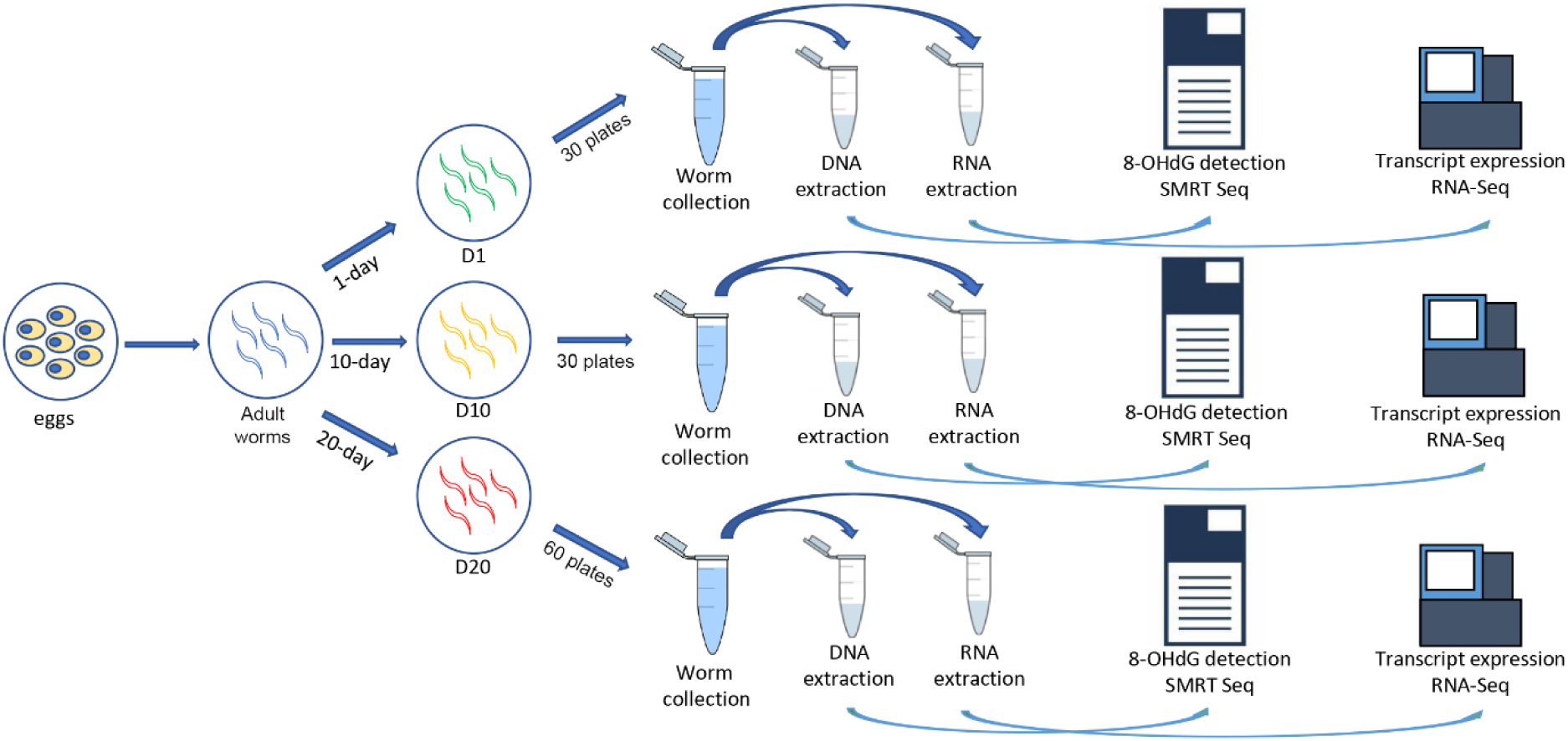
The workflow of *C. elegans* sample preparation. In brief, the adult worms were cultured into indicated number of plates until the required time periods and collected for further experiments. To maintain the cultured environment and worm batch between DNA and RNA samples consistent, the collected worms were subdivided into two tubes for each DNA and RNA preparation. (n=30 plates for D1, n=30 plates for D10, and n=60 plates for D20, respectively.)

**Figure S5.**
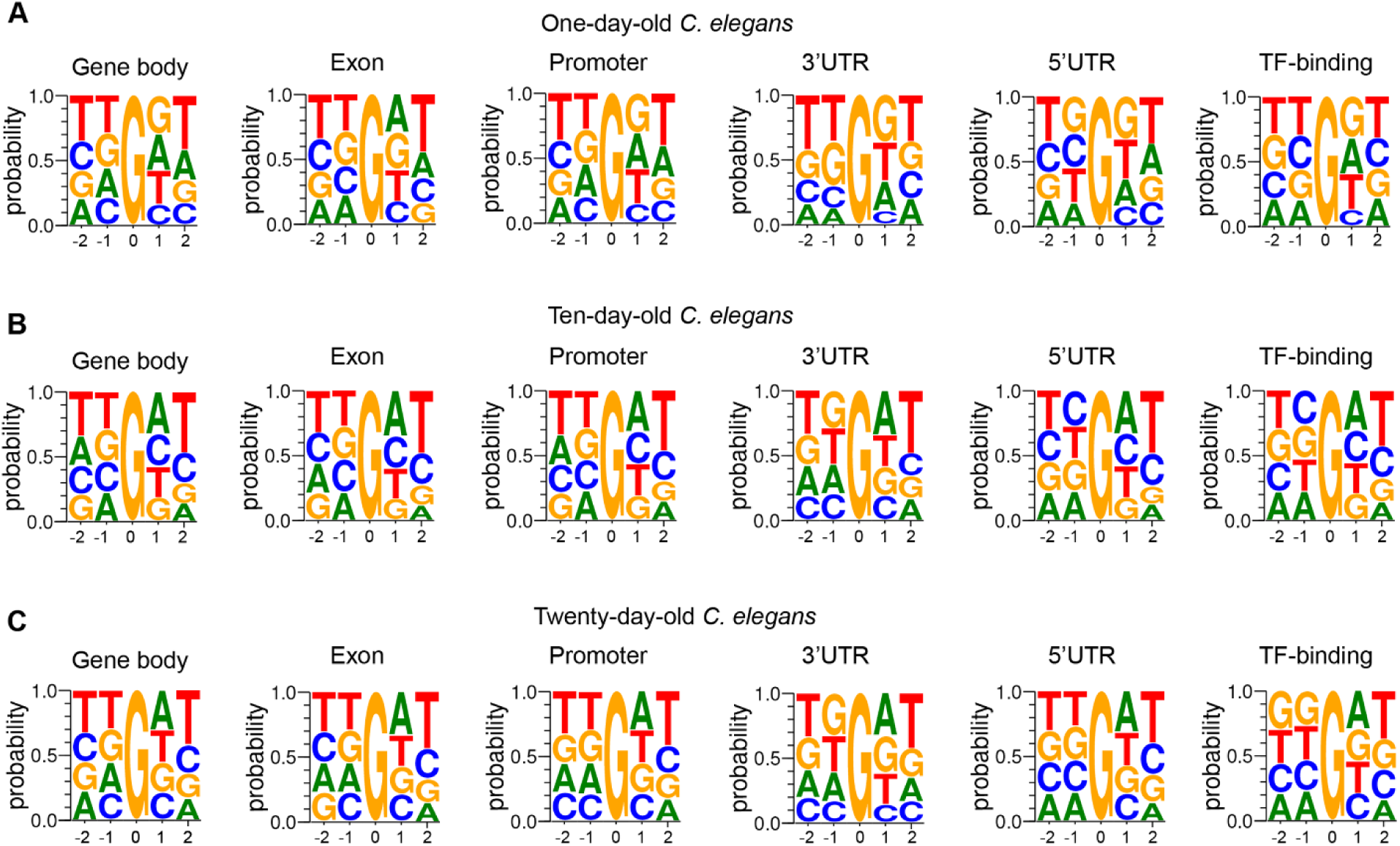
Consensus sequence next to 8-OHdG detected in different genomic features of *C. elegans.* **A-C.** Sequence logo plots show the local sequence context (−2bp and +2bp) surrounding to 8-OHdG in different genomic features of different age groups: 1-day-old *C. elegans* (A), 10-day-old *C. elegans* (B); and 20-day-old *C. elegans* (C).

**Figure S6.**
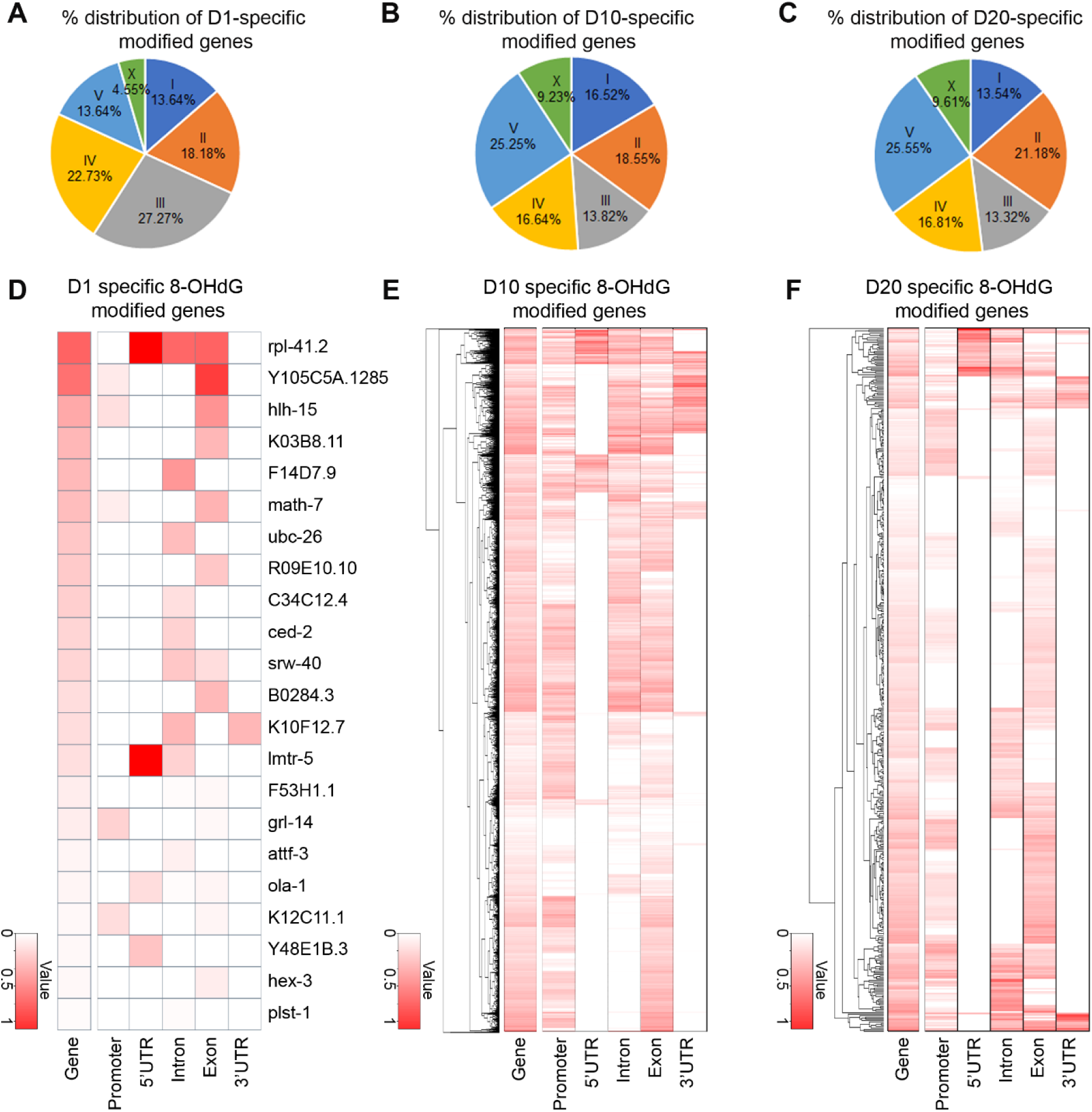
Distribution of age-specific oxidatively modified genes. **A-C**. The % distribution of age-specific 8-OHdG modified genes across different chromosomes. **D-F**. Heatmap of 8-OHdG modification process in different genomic regions. Genes exclusively oxidized in 1-day-old *C. elegans* (A, and D), those exclusively oxidized in 10-day-old *C. elegans* (B, and E); those exclusively oxidized in 20-day-old *C. elegans* (C, and E). The 8-OHdG modification sites in the whole length of the encoded gene were counted using customized Perl scripts and we assigned these genes as age-specific only if the 8-OHdG sites in the gene region of exclusively in one sample but not in the other sample, with a significant *p*-value < 0.05 given by multiple *Fisher* exact test and a *Benjamini–Hochberg* false discovery rate (FDR) < 0.05. The differential oxidation coordinates were identified by the number of 8-OHdG sites in 100 kb-length within the indicated regions. The color key from white to red indicates 8-OHdG level from low to high, respectively.

**Figure S7.**
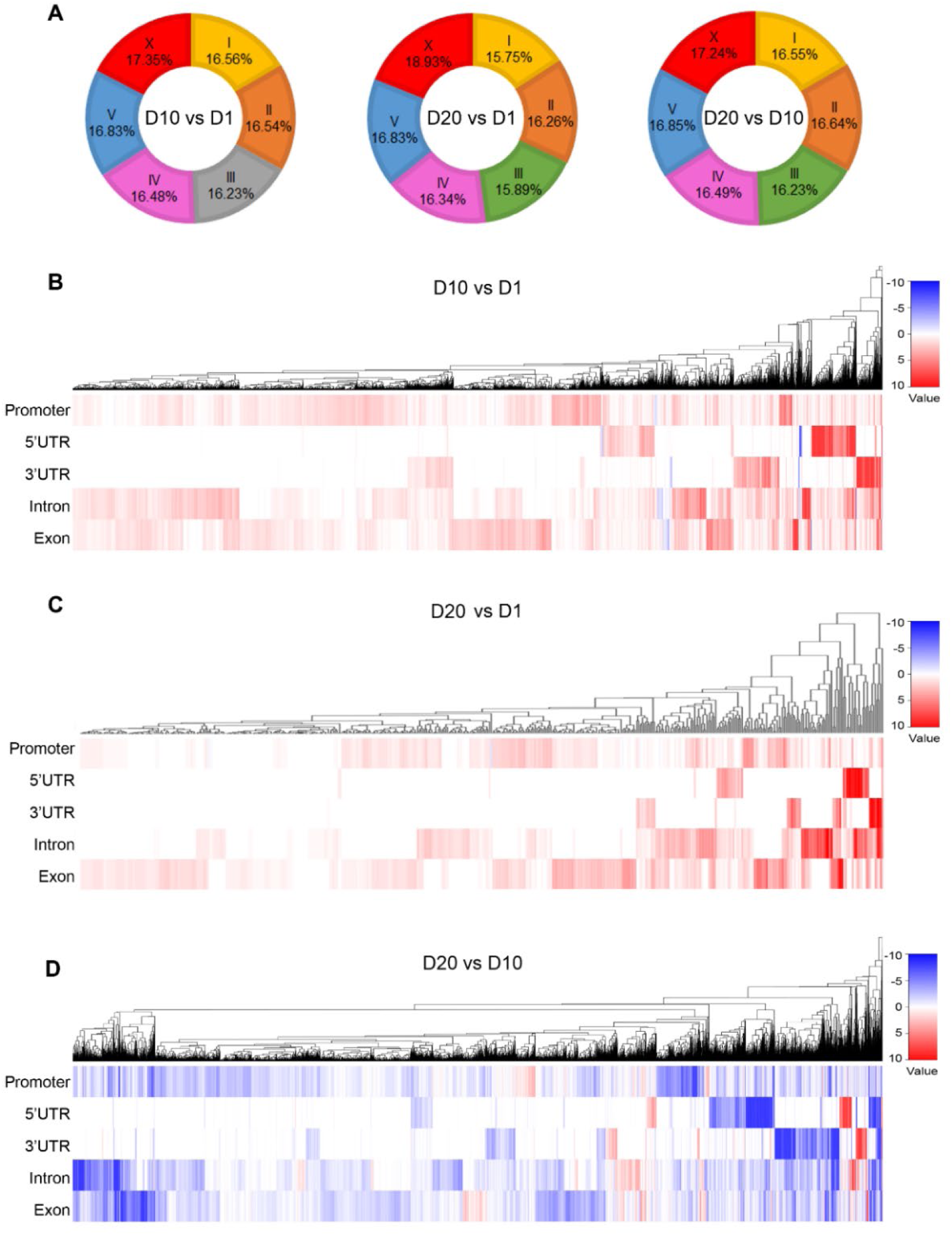
Genomic distribution of differential 8-OHdG modified genes. **A**. The proportion of differential 8-OHdG modified genes (DOGs) captured in each chromosome. DOGs were identified by comparing the number of 8-OHdG sites observed in the same gene annotated between two samples. The 8-OHdG modification sites in the whole length of the encoded gene were counted using customized Perl scripts and we assigned these genes as DOGs only if the 8-OHdG sites in the gene region of one sample was greater than or less than that of another sample, with a significant *p*-value < 0.05 given by multiple *Fisher* exact test and a *Benjamini–Hochberg* false discovery rate (FDR) < 0.05. **B-D**. Heatmap of differential 8-OHdG modification process in different genomic regions analyzed between D10 vs D1 (B), between D20 vs D1 (C), and between D20 vs D10 (D). The differential oxidation coordinates were identified by the number of 8-OHdG in 100 kb-length within the indicated regions with a *Benjamini–Hochberg* false discovery rate (FDR) < 0.05. The color key from blue to red indicates 8-OHdG level from low to high, respectively.

**Figure S8.**
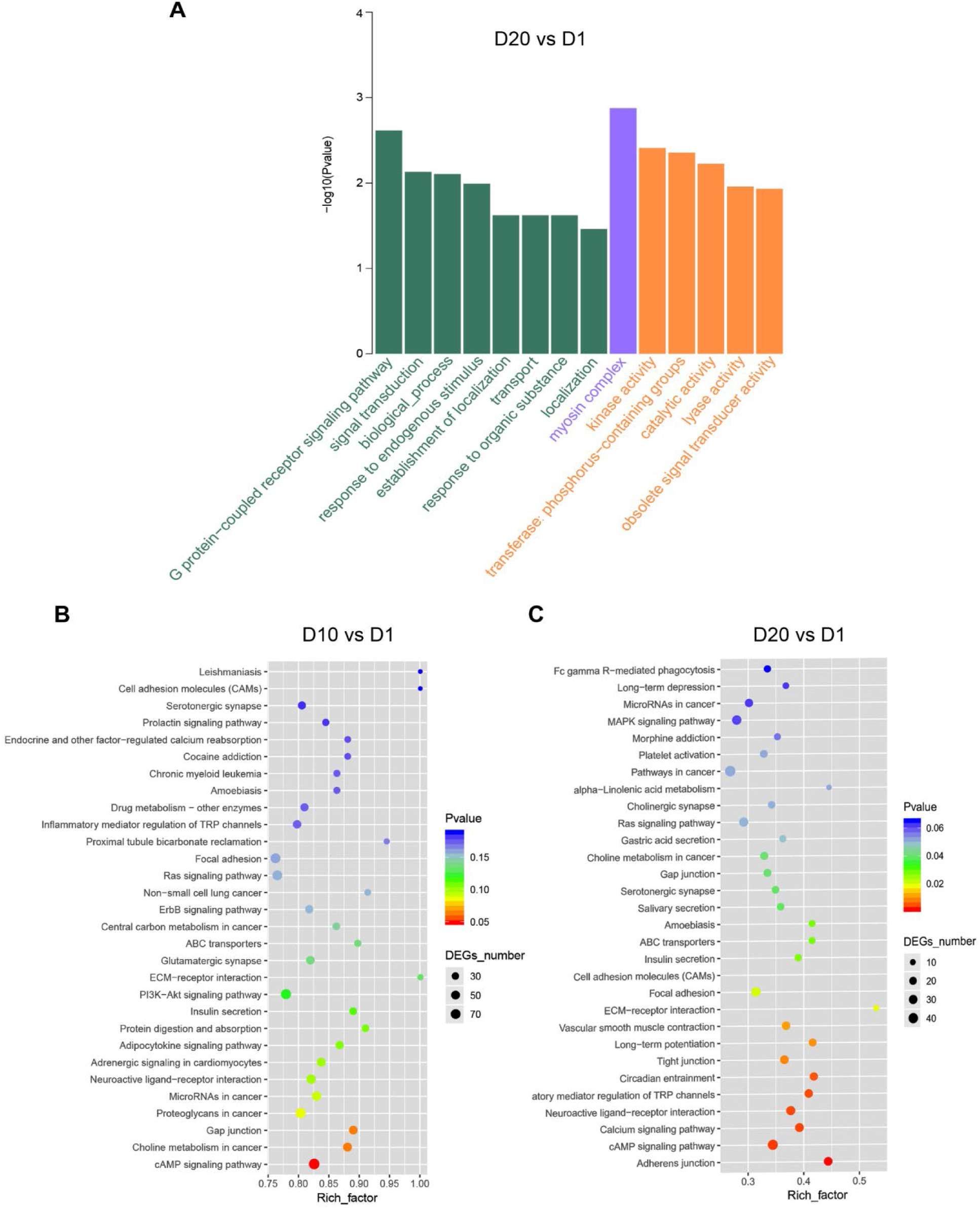
Functional annotations for differentially 8-OHdG modified genes (DOGs). GO (Gene ontology, A) and KEGG (Kyoto Encyclopedia of Genes and Genomes, B and C) analysis for DOGs (fold change modification bases in each gene between D10 *vs.* D1 or D20 *vs.* D1).

**Figure S9.**
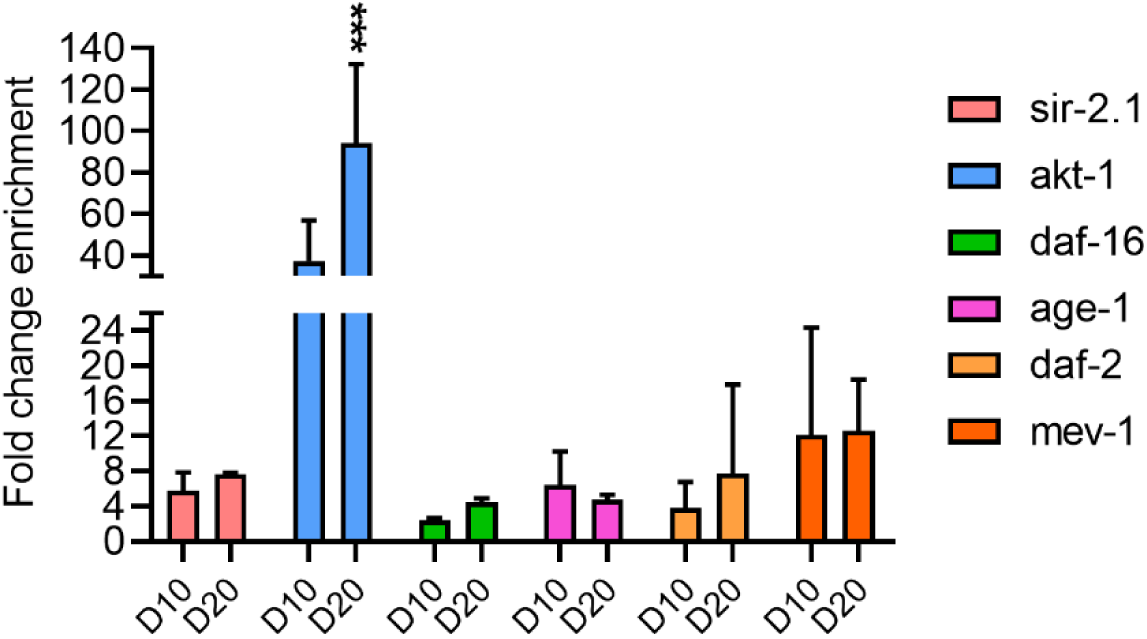
8-OHdG modification in 6 longevity regulating genes. Validation of the 6 identified 8-OHdG modified longevity regulating genes loci from SMRT sequencing by 8-OHdG-IP-qPCR assay (n=3). *Act-1* was used as a negative control. Fold change enrichment is identified by comparing with expression level in D1 samples. Results are represented as mean ± SEM. \*\*\**P*< 0.005 compared to D10.

**Figure S10.**
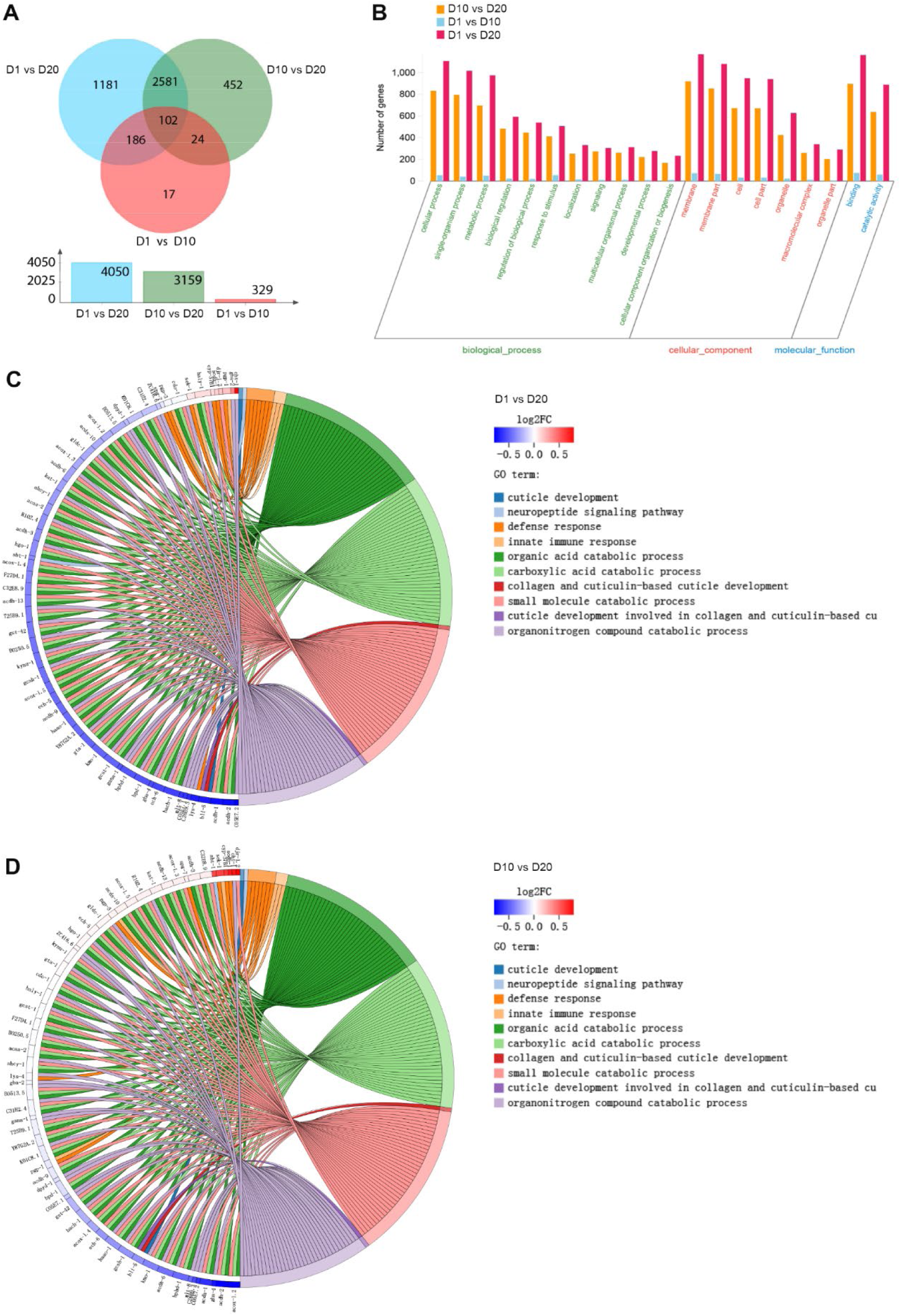
The age-specific pattern of transcript expression in *C. elegans*. **A**. Venn diagram comparison of transcript expressions in different samples. **B**. The GO pathway analysis of differentially transcribed genes among different samples. **C and D**. KEGG analysis of differentially transcribed genes between D1 *vs.* D20 (C) and D10 *vs.* D20 (D).

**Figure S11.**
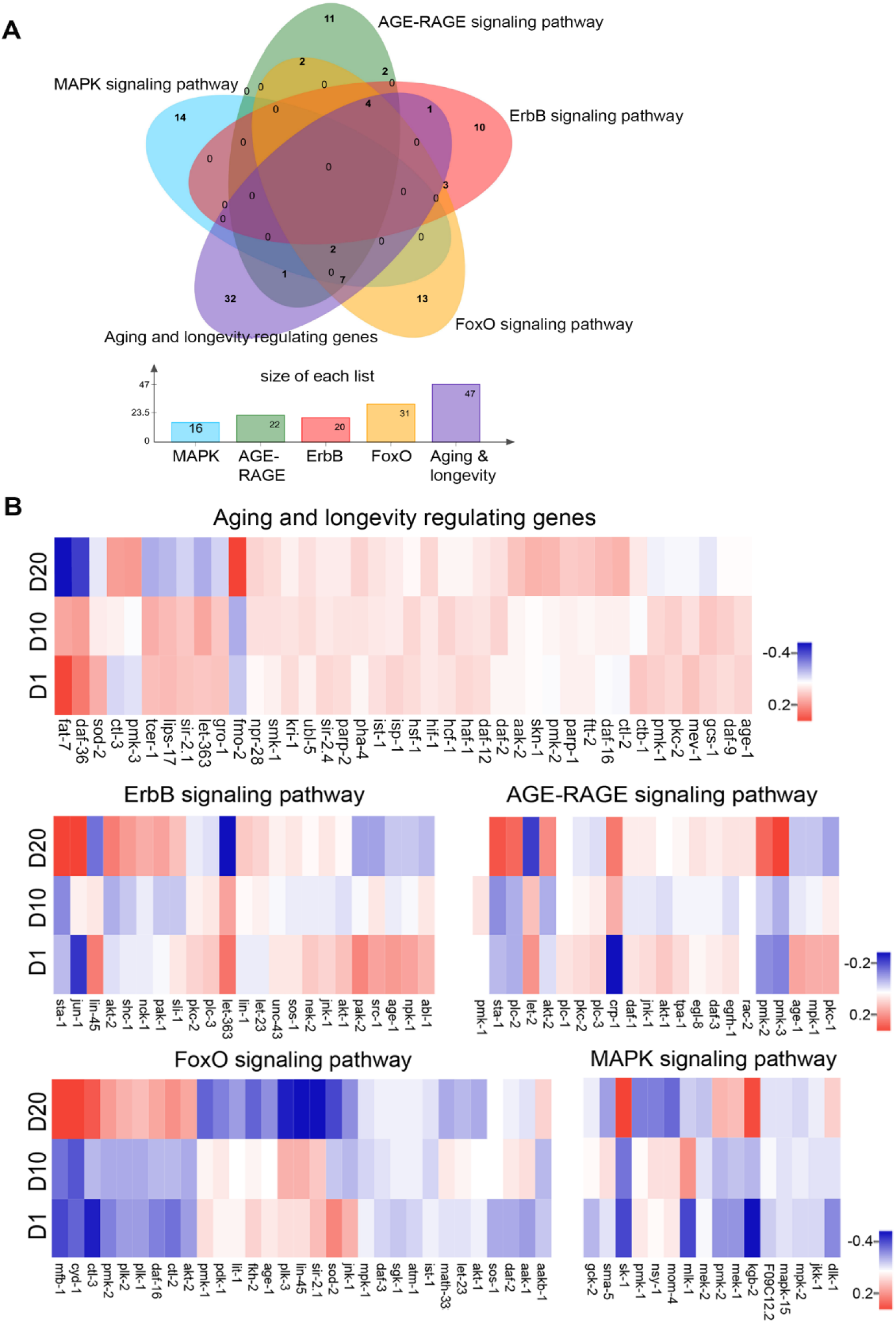
The potential correlation between 8-OHdG modification and transcript dynamics of age-regulating genes. **A**. Venn diagram comparison of transcript expression levels for genes involved in key pathways enriched by DOG (Fig. 3F). **B**. Heatmap of transcript expression levels of DOG involved in top-5 pathways. The log2 of the transcript expression levels (RPKM) of RefSeq genes were calculated as described (Trapnell et al., 2012). The color key from blue to red indicates the expression levels (RPKM) low to high, respectively.

**Figure S12.**
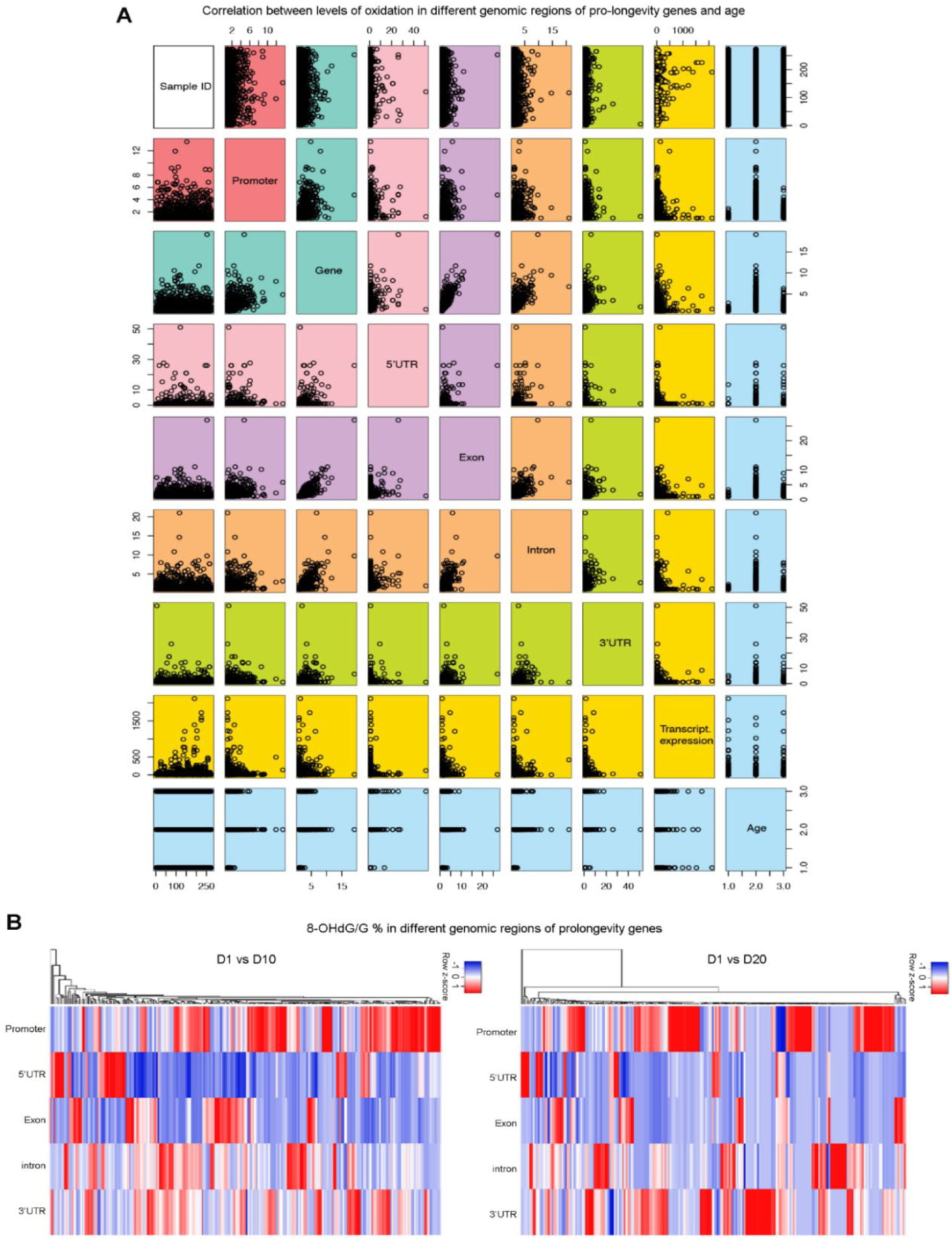
Correlation analysis of 8-OHdG modification and transcript expression of pro-longevity genes. **A**. Pair-wise correlation analysis for 8-OHdG modification across different genomic regions and transcript expression of pro-longevity genes with aging. The log2 of the transcript expression levels (RPKM) of RefSeq genes were calculated as described (Trapnell et al., 2012). The scatter-plot matrix is shown on the side of the diagonal. **B**. Heatmap showing the genomic distribution of 8-OHdG in pro-longevity genes. The color key from blue to red indicates 8-OHdG level from low to high, respectively.

## Supplementary Tables

**Table S1:** Summary of sequencing samples.

| Samples | Genome size | Library | Coverage per strand | Subread N50 | Total Data (Gb) |
| --- | --- | --- | --- | --- | --- |
| Synthetic oligonucleotide sequence | 155 bp | 155 bp | ~340x | 953 | 0.4 |
| Plasmid control | 5.4 kbp | ~5 kb | 1000x | 3,474 | 5.6 |
| Plasmid treated with H<sub>2</sub>O<sub>2</sub> | 5.4 kbp | ~5 kb | 1000x | 1,577 | 0.3 |
| Plasmid treated with H<sub>2</sub>O<sub>2</sub> and Ogg1 | 5.4 kbp | ~5 kb | 1000x | 1,387 | 5.5 |
| Plasmid (G2427T mutation) treated with H<sub>2</sub>O<sub>2</sub> | 5.4 kbp | ~5 kb | 1000x | 3,403 | 8.4 |
| Plasmid (random mutation) treated with H<sub>2</sub>O<sub>2</sub> | 5.4 kbp | ~5 kb | 1000x | 3,403 | 2.3 |
| 1-day-old *C. elegans* | 102 Mbp | 10 kb | 104x | 7,860 | 16.3 |
| 10-day-old *C. elegans* | 102 Mbp | 10 kb | 134x | 9,580 | 28.9 |
| 20-day-old *C. elegans* | 102 Mbp | 10 kb | 100x | 9,540 | 22.9 |

Summary of samples used in this study. The coverage for each sample was determined by the mean reference genome alignment depth. All subread coverage were estimated using raw sequence data throughput and a genome size of 102 Mbp. Synthetic oligonucleotide sequence was sequenced with P6-C4 chemistry in RSII platform, and all remaining samples were sequenced with v2.1 chemistry in Sequel platform. 10 μM of H<sub>2</sub>O<sub>2</sub> was used for oxidative stress induction; 10 μg of Ogg1 was used for base excision reaction.

**Table S2: Motif analysis of 8-OHdG sites in different age groups.** (Related to Figure 1)

**Table S3: Summary of 8-OHdG modified genes in the *C. elegans* Genome.** (Related to Figure 3)

**Table S4: Relative frequency of 8-OHdG modification identified in the different genomic regions of one-day-old *C. elegans*.** (Related to Figure 3 and S6)

**Table S5: Relative frequency of 8-OHdG modification identified in the different genomic regions of ten-day-old *C. elegans*.** (Related to Figure 3 and S6)

**Table S6: Relative frequency of 8-OHdG modification identified in the different genomic regions of twenty-day-old *C. elegans*.** (Related to Figure 3 and S6)

**Table S7: Relative frequency of 8-OHdG modification identified in the different genomic regions of longevity regulating genes.** (Related to Figure 5)

**Table S8: The Primers used for 8-OHdG-IP-qPCR.** (Related to Figure S9 and supplementary methods)

